# Polyjet 3D printing of tissue-mimicking materials: how well can 3D printed synthetic myocardium replicate mechanical properties of organic myocardium?

**DOI:** 10.1101/825794

**Authors:** Leah Severseike, Vania Lee, Taycia Brandon, Chris Bakken, Varun Bhatia

## Abstract

Anatomical 3-D printing has potential for many uses in education, research and development, implant training, and procedure planning. Conventionally, the material properties of 3D printed anatomical models have often been similar only in form and not in mechanical response compared to biological tissue. The new Digital Anatomy material from Stratasys utilizes composite printed materials to more closely mimic the mechanical properties of tissue. Work was done to evaluate Digital Anatomy myocardium under axial loading for comparison with porcine myocardium regarding puncture, compliance, suturing, and cutting performance.

In general, the Digital Anatomy myocardium showed promising comparisons to porcine myocardium. For compliance testing, the Digital Anatomy was either within the same range as the porcine myocardium or stiffer. Specifically, for use conditions involving higher stress concentrations or smaller displacements, Digital Anatomy was stiffer. Digital Anatomy did not perform as strongly as porcine myocardium when evaluating suture and cutting properties. The suture tore through the printed material more easily and had higher friction forces both during needle insertion and cutting. Despite these differences, the new Digital Anatomy myocardium material was much closer to the compliance of real tissue than other 3D printed materials. Furthermore, unlike biological tissue, Digital Anatomy provided repeatability of results. For tests such as cyclic compression, the material showed less than two percent variation in results between trials and between parts, resulting in lower variability than tissue. Despite some limitations, the myocardium Digital Anatomy material can be used to configure structures with similar mechanical properties to porcine myocardium in a repeatable manner, making this a valuable research tool.

## Introduction

Anatomical 3-D printing has the potential to be used for various applications related to visualization and education, research and development processes, implant training, and pre-procedure planning. Anatomical 3-D printing also has the potential to bridge the testing workflow between ex vivo benchtop testing and in vivo animal testing with a lower barrier to access; however, commercially available 3D printing materials are limited to relatively rigid materials. Additionally, there is limited quantitative comparison of mechanical properties of these material to organic tissue mechanical properties. Some research exists for heart valves and bone comparisons, but not for myocardium [1, 2]. The work reported in this paper aims to quantify and compare application specific mechanical properties of Digital Anatomy 3D printing materials to equivalent porcine tissue properties.

Stratasys (Eden Prairie, MN) has developed a suite of new Digital Anatomy materials designed to mimic anatomical tissues, including materials to mimic myocardial tissue. In general, these materials have varying degrees of compliance, and are printed as a shell of Agilus material (Hardness = 30A) filled with a mix of TissueMatrix (Hardness = 00A) and Agilus in a gyroid structure infill pattern, which is varied to change compliance.

Porcine myocardium was chosen as the baseline for comparison because of the similarity to human tissue, availability, and precedent for preclinical testing of cardiac devices.

The tests encompassed in the work included tensile testing, compliance testing, puncture testing, suture testing, and qualitative cutting comparisons. These tests allowed quantification of the printed myocardium and an understanding of how they compare to the porcine myocardium.

## Methods

### Material options for printing

For printing myocardium, Digital Anatomy offers five material options. Throughout the testing process, some materials changed either in name or infill formula. Table 1 below maps the name and material changes and details the naming conventions that will be used throughout the paper.

**Table 1:** Naming convention for the 3D printed materials throughout the paper, commercial release names, and material descriptions.

| Paper Name | Release name | Material description |
| --- | --- | --- |
| Myocardium A* | N/A | Pure TissueMatrix wrapped in 0.4 mm Agilus |
| Myocardium 1 | Highly Contractile | Softest infill wrapped in 0.4 mm Agilus |
| Myocardium 2 | Moderately Stiff | Second softest infill wrapped in 0.4 mm Agilus |
| Myocardium 3 | Stiffened | Second stiffest infill wrapped in 0.4 mm Agilus |
| Myocardium 4 | Very stiff | Stiffest infill wrapped in 0.4 mm Agilus |
| Myocardium 5 | Extremely Stiff | Softest infill wrapped in 0.6 mm Agilus |
| Myocardium B* | N/A | Pure TissueMatrix wrapped in 0.6 mm Agilus |

### Printing and Cleaning

Samples were batch printed exclusively for the outlined testing using the recommended support structure. Cleaning was done with water using a pressurized washer. No additional chemicals were used in the cleaning process, and testing was completed within a week of printing.

### Tensile Testing

Tensile testing was performed according to ASTM Standard 638. Type IV dog bones were selected with dimensions shown in Figure 1. While this standard is normally reserved for homogenous materials, this test was used to quantitatively compare the composite 3D printed materials described in Table 1. In order to prevent failure at the pneumatic grips, the samples were printed with mechanically interlocked rigid grip sections to accommodate softer durometer materials, as shown in Figure 2. Six samples were tested of each material type.

**Figure 1:**
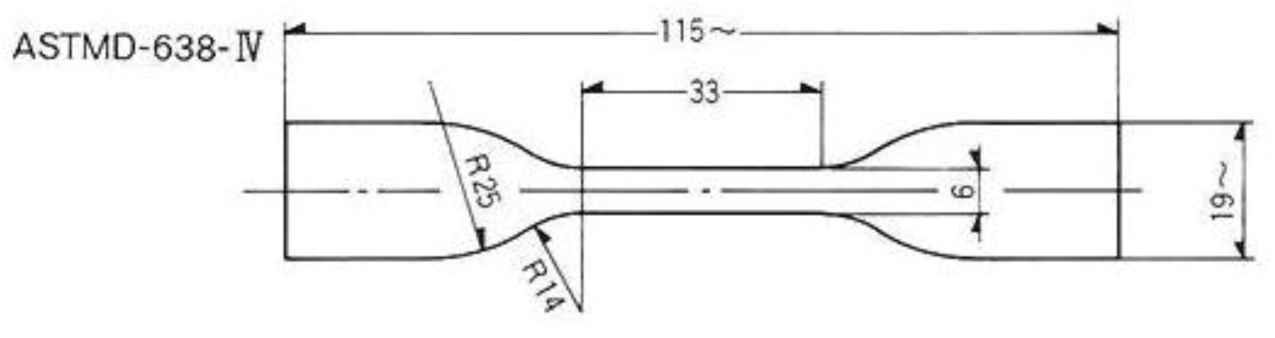
ASTM 638-V Type IV dog bone dimensions in millimeters

**Figure 2:**
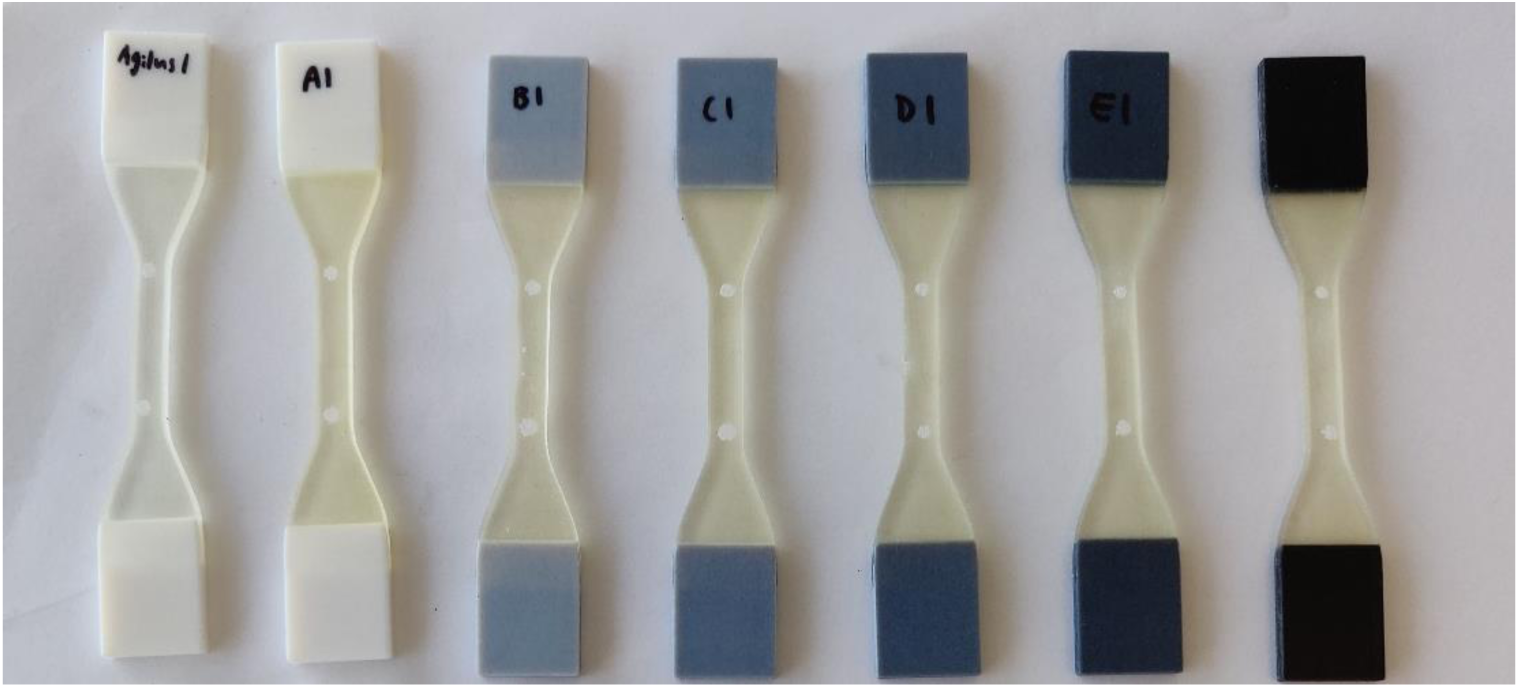
Printed dog bones with Vero grips. Left to right is Agilus, Myocardium A*, Myocardium 1, Myocardium 2, Myocardium 3, Myocardium 4, Myocardium 5, and Myocardium B*.

Testing was performed on a dual column Instron with a Digital Image Correlation (DIC) camera for tracking extension. The parts were held on the Instron with 80 PSI pneumatic grips. Stress and strain measurements were obtained from this test, and an analogue to Young’s Modulus was calculated.

### Tissue Collection

Porcine hearts were obtained from two different sources, a butcher and a pre-clinical research facility, in either a frozen or recently harvested unfrozen state. Frozen porcine hearts were purchased on August 16, 2019 from a butcher shop and immediately moved to a −80C freezer [3]. According to the supplier, the hearts were harvested on July 8, 2019 and kept frozen until the purchase date. Additional porcine hearts were harvested from termed pre-clinical research animals. Hearts were excised within two hours of termination, dissected, and samples were frozen in a −80C freezer within four hours of termination. Alternatively, hearts were frozen in a −80C freezer immediately following removal and thawed and dissected on the day of testing. Frozen hearts were wrapped in saline soaked paper towels.

For sample preparation, hearts were thawed incrementally to dissect and test without the need to refreeze the samples. Hearts were thawed by placing frozen hearts in an insulated cooler overnight. Hearts were checked incrementally starting at about 14 hours of thaw time. Once hearts were thawed enough to cut, they were dissected in order to isolate the tissue from the different chambers. Samples were excised in a two-inch circle and then transferred into saline until it was time to test. Prior to testing, samples were blotted dry using paper towels. Samples were tested within hours of dissection.

The porcine hearts were dissected to isolate the different portions of the heart as shown in Figure 3: right atrium (RA), left atrium (LA), left ventricle (LV), right ventricle (RV), and ventricular septum (VS). As shown in Figure 4, a two-inch diameter circle of tissue was cut using a punch and prepared for testing. Samples were taken in order to maximize the number of samples per heart, and tested with the probe travelling from endocardium to epicardium. For the ventricles, samples were taken from the free wall. For the atria, most samples were taken from the appendages due to the small size of the free walls. When possible, atrial samples were also taken from the free wall. Samples from the ventricular septum were tested from the right side. Samples from the atrial septum could not be obtained because of the small size.

**Figure 3:**
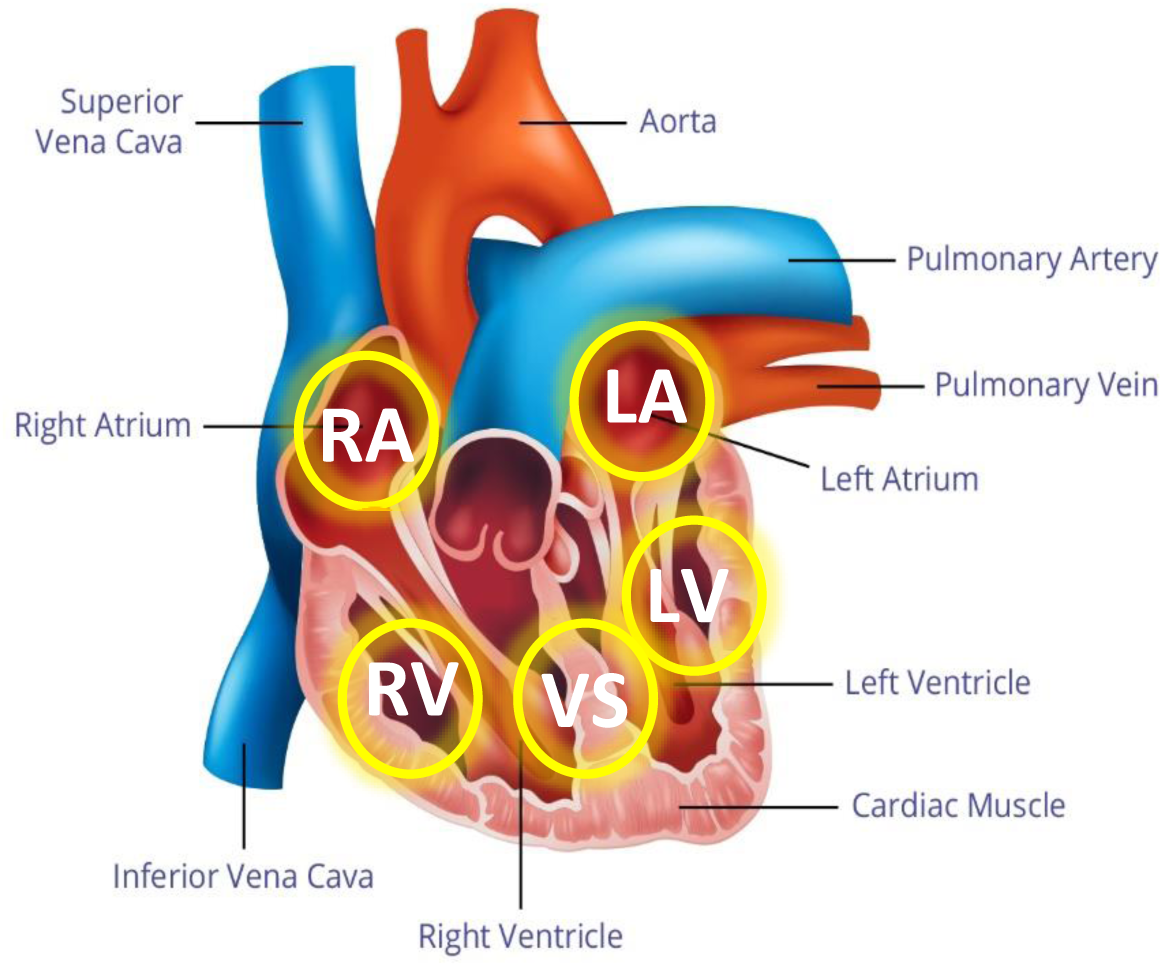
Where the tissue samples were taken from the heart. Right atrium (RA), left atrium (LA), right ventricle (RV), left ventricle (LV), and ventricular

**Figure 4:**
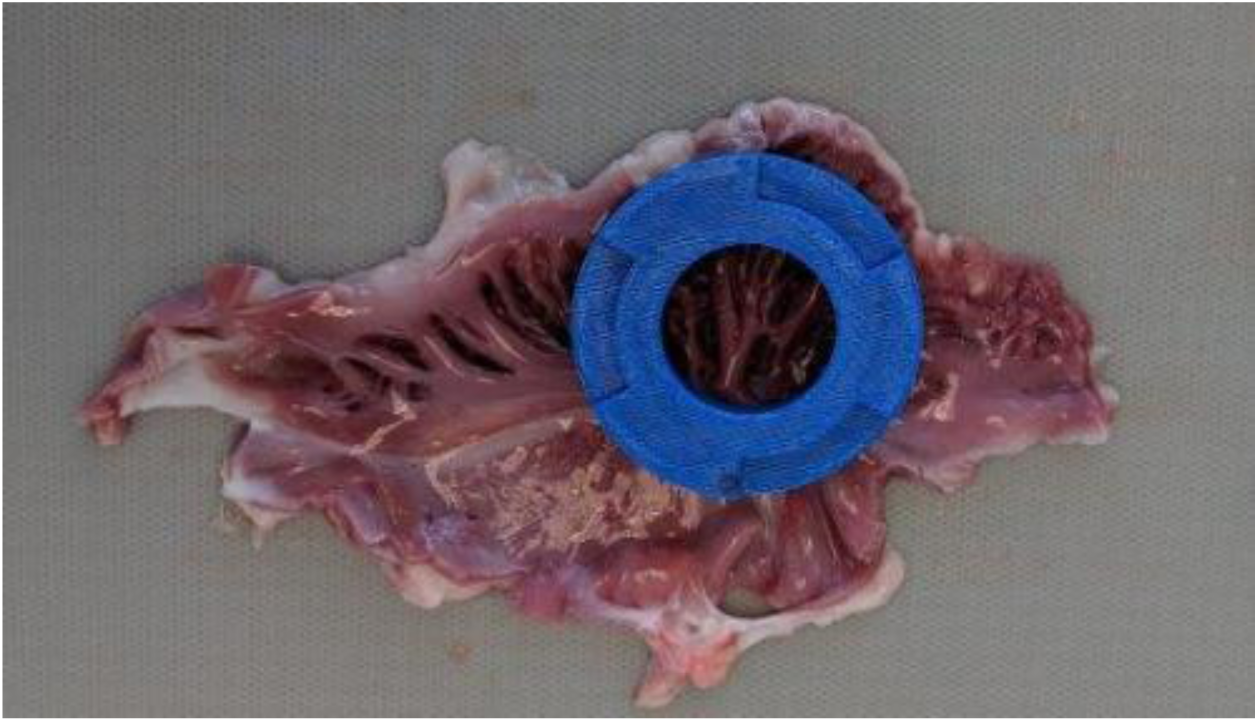
An example of where samples were cut out of the right atrium. In this case, the sample was cut out of the right atrial appendage.

### Puncture Testing

Puncture testing was performed on two-inch diameter tissue samples clamped over a one inch diameter hole. These dimensions were chosen to better mimic the boundary conditions found in the heart. The test fixture was designed to sit centered on the bottom of an Instron, with the clamping piece elevated to allow for tenting. The samples were then clamped between the fixture and a top plate. Testing was completed using a 50 N load cell at a compression rate of 0.5 in/min; the setup is shown in Figure 5.

**Figure 5:**
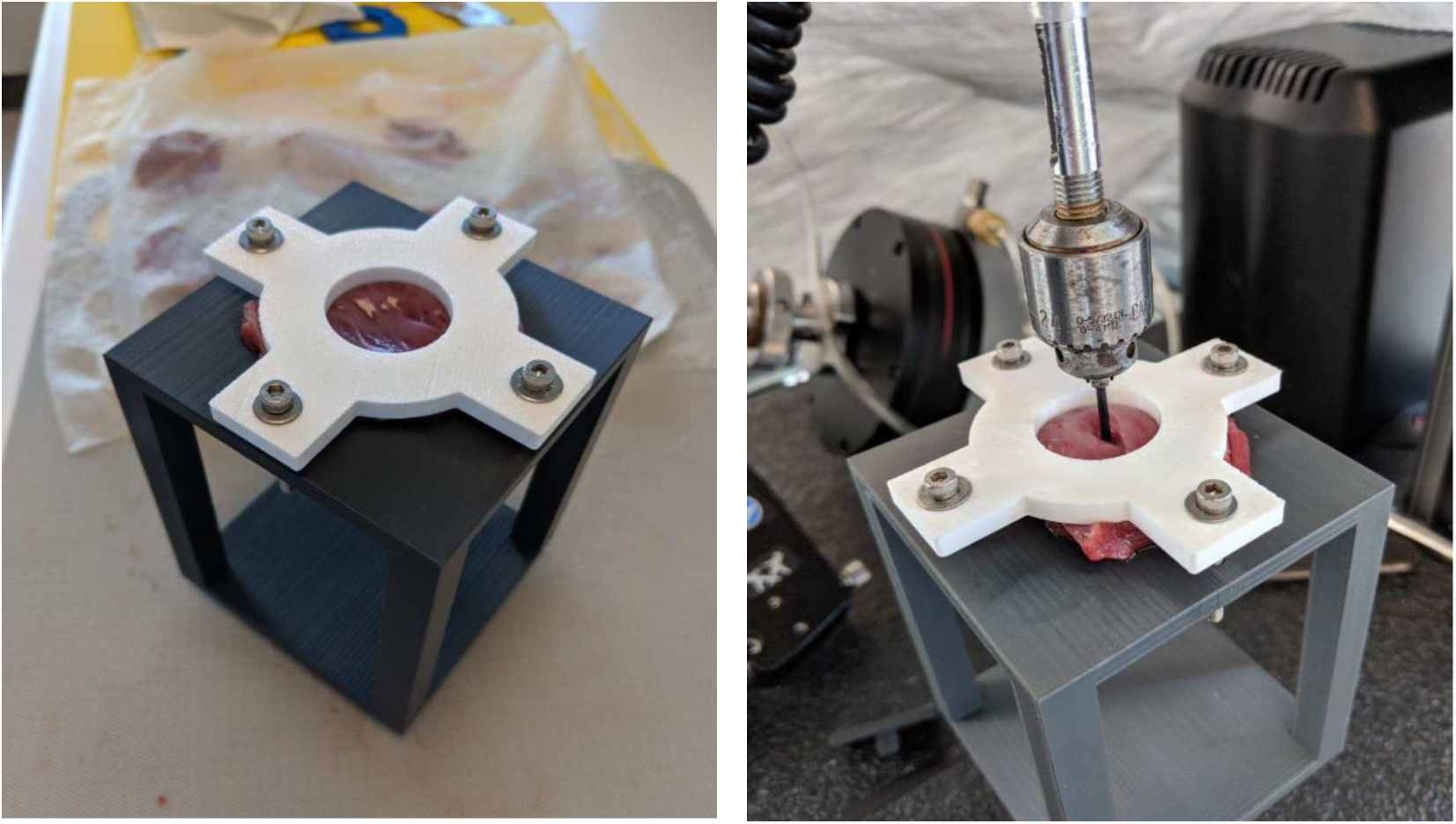
The test fixture on the left. The sample is clamped between an elevated plate with a hole and a top plate using bolts. On the right, the fixture and sample is in use on the Instron, with a constrained pin attached to the load cell.

Screening tests were performed to find the printed materials and test methods that were most relevant for further testing. A few parameters were evaluated including material type, hole diameter, material thickness, and rod diameter. The combinations which were of interest are listed in Table 2. Two samples of each combination were tested to help narrow down material types for further testing. Some material changes occurred during testing, so not all materials were tested at every combination.

**Table 2:** Variable parameters used for filter testing for the puncture and compliance test.

| Parameter | Options |
| --- | --- |
| Material | Agilus30<br>Myocardium 1<br>Myocardium 2<br>Myocardium 3<br>Myocardium 4<br>Myocardium 5 (only partially tested)<br>Myocardium A* (only partially tested)<br>Myocardium B* (only partially tested) |
| Thickness | 5 mm (roughly corresponds to right ventricular thickness)<br>20 mm (roughly corresponds to left ventricular thickness) |
| Rod diameter | 9 French (3 mm)<br>22 French (7.3 mm) |
| Hole diameter | One inch<br>Two inch |

Printed thicknesses were designed based on literature values corresponding to porcine hearts. For healthy hearts, average left ventricular thickness was listed as 20 mm and right ventricular thickness was listed as 5 mm [4]. Puncture pin diameters were chosen based on typical cardiac use cases in the cardiac rhythm space such as a traditional pacing system - a battery implanted in the shoulder region connected to wires, or leads, that are fixated at various locations in the heart. To represent smaller cardiac lead delivery systems, a 9 French pin (3 mm diameter) was used. To represent larger device delivery systems, such as Medtronic’s leadless pacing system, a 22 French pin (7.3 mm diameter) was used.

The printed sample parameters with the most diverse results were further tested to sample sizes of six to quantify their mechanical properties, and their comparison to different sections of the porcine myocardium. The parameters listed in Table 3 describe the samples that were tested further.

**Table 3:** Final testing parameters for puncture and compliance testing of printed disks.

| Parameter | Options |
| --- | --- |
| Material | Myocardium 1<br>Myocardium 4<br>Myocardium 5 |
| Thickness | 2.7 mm<br>6.7 mm<br>12.1 mm |
| Rod diameter | 9 French<br>22 French |
| Hole diameter | One inch |

### Compliance Testing

The compliance testing analyzed up to the first 10 mm beyond the pre-load from puncture testing, a relevant displacement for typical use cases. A pre-load of 1 N was chosen to minimize the noise of the tissue data due to trabeculae and other structures, as shown in Figure 6. Starting at 1 N, the slope of best fit was taken for 0-5 mm of displacement and 5-10 mm of displacement. For printed material, the first peak force corresponds to the first punctured Agilus layer. In the tissue samples, this would be the endocardial tissue layer depending on test orientation.

**Figure 6:**
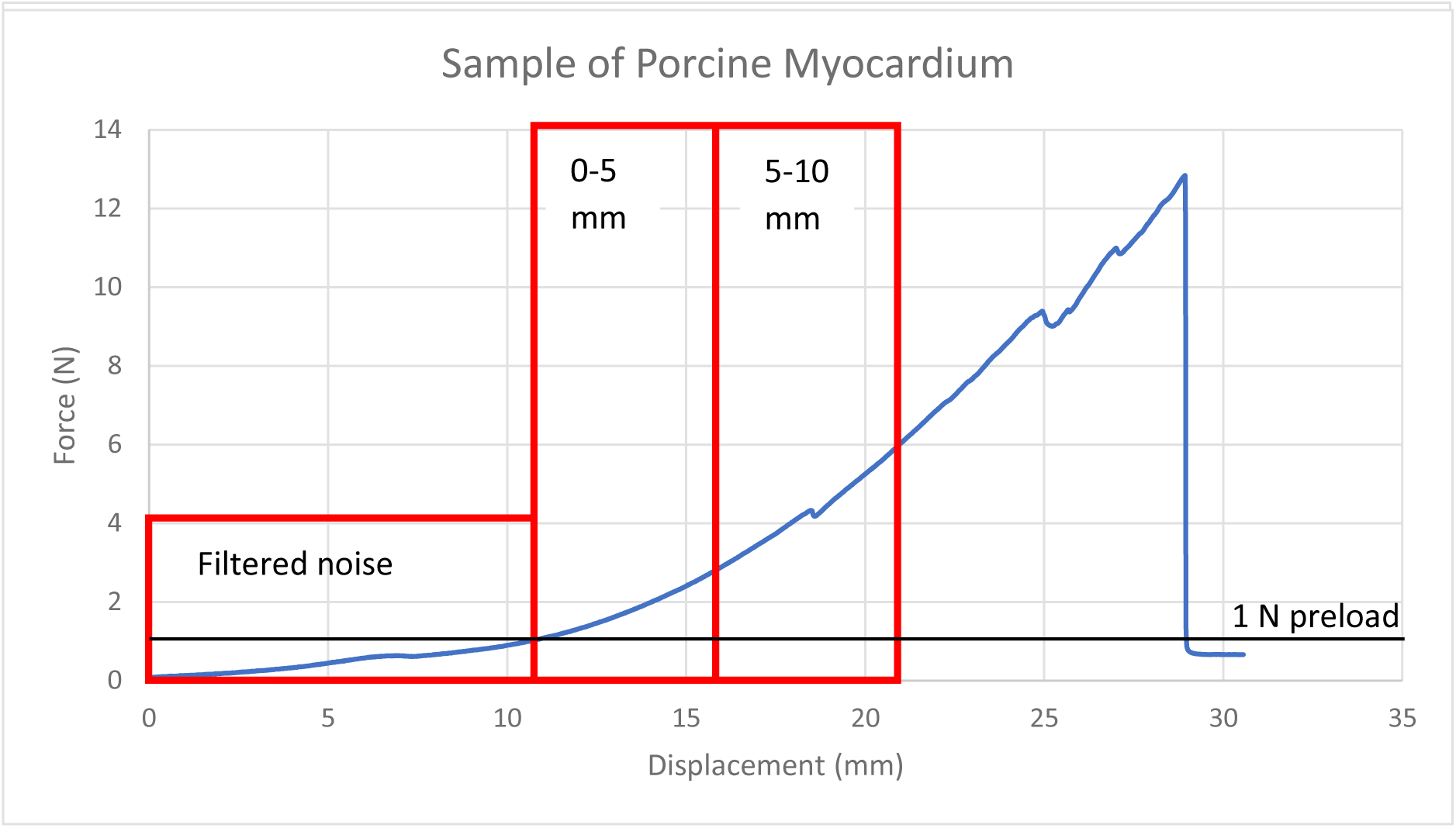
An example of a porcine myocardium puncture test. Regions of interest are the noise that was filtered out, and the two levels of displacement analyzed for compliance.

### Compliance Repeatability Testing

Additional testing was performed to assess the repeatability of use of the printed material using the subset of parameters listed in Table 4. Four samples of each combination were tested using the same fixture and load cell as the puncture and compliance test. The pin was displaced to 0.1 N of pre-load, extension zeroed, and then displaced 10 mm into the samples. The pin was then returned to the starting position, idle for 10 seconds, and displaced 10 mm again for a total of five cycles. The data was then analyzed using the same methods as compliance testing. Starting at 1 N of force, a slope was fit for 5 mm of displacement. The percent difference in slope was calculated between the first curve and the four following curves.

**Table 4:**
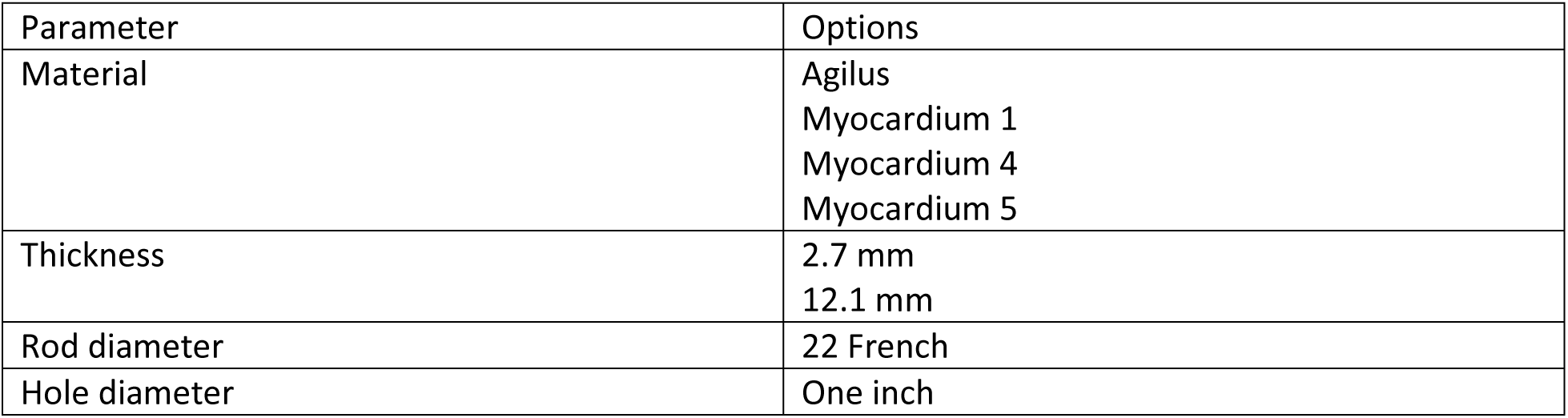
Parameters for compliance repeatability testing of printed disks.

### Suture Testing

Suture testing was done on printed materials to compare retention force to that of porcine tissue. Five left ventricle samples and two right ventricle samples were excised from 2 porcine hearts and tested against two samples each of Agilus, Myocardium 1, and Myocardium 5 at 12.1 mm and 2.5 mm thicknesses.

The test setup used the same fixture set up as the puncture and compliance testing, but instead of compressive force, extension force was measured. EthiBond Excel Green Braided Polyester 3-0 17 mm 1/2c Taper suture was used on all samples, and suture stiches were performed by a trained veterinary scientist. Suturing using normal tie down force resulted in prematurely torn samples. As shown in Figure 7, the sutures were intentionally kept loose to prevent tearing of the printed material prior to testing, and the same suture technique was replicated on the porcine myocardium for appropriate comparison. This stitch was then looped around a hook on an Instron and pulled at 0.5 inches/minute until the suture was free of the sample. Peak forces for each sample were recorded.

**Figure 7:**
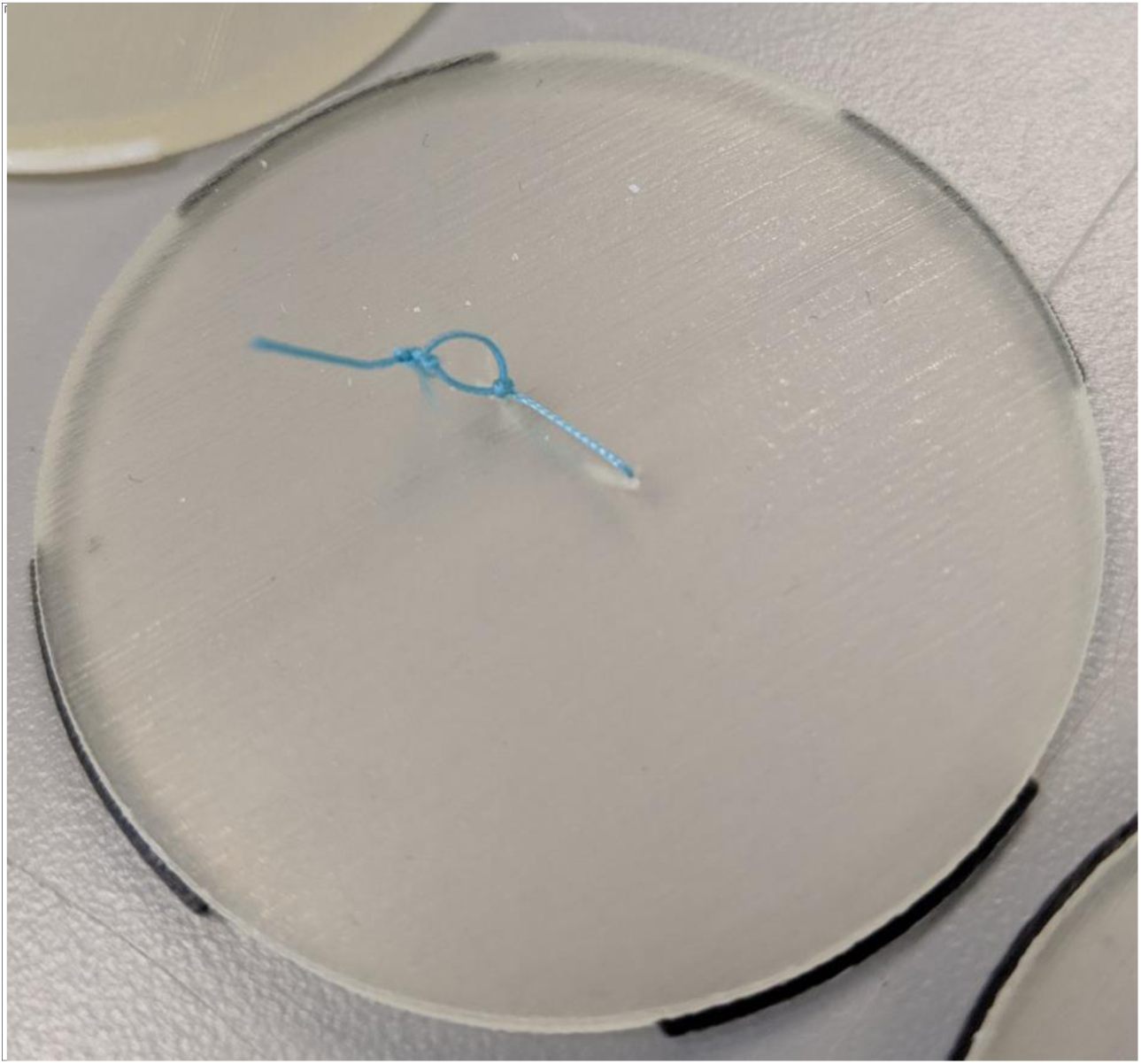
A single suture on a 2.5 mm thick disk of Myocardium 5. Note how loose the suture is tied.

### Qualitative Suture and Cutting Force

Four expert reviewers were asked to suture and cut samples of 12.1 mm thick Agilus, Myocardium 1, and Myocardium 5 and compare the experience to animal and cadaver myocardium. Reviewers were asked to evaluate the similarity between the printed materials and porcine hearts, past animal experience, and past cadaver experience on a scale of 1 - 7. Comments were also provided by the reviewers.

## Results and Discussion

### Tensile Testing

The analogue for Young’s Modulus was determined using a linear fit up to 50% strain, as shown in Table 5. All materials had a sample size of six except for Agilus, which had five.

**Table 5:** Analogue of Young’s Modulus Values for printed materials.

| Material | Modulus (N/m<sup>2</sup>) |
| --- | --- |
| Agilus | $0.536 \pm 0.009$ |
| Myocardium A* | $0.262 \pm 0.004$ |
| Myocardium 1 | $0.294 \pm 0.003$ |
| Myocardium 2 | $0.310 \pm 0.003$ |
| Myocardium 3 | $0.327 \pm 0.005$ |
| Myocardium 4 | $0.342 \pm 0.007$ |
| Myocardium B* | $0.334 \pm 0.010$ |

The moduli of the printed materials ranged from 0.262 to 0.536 MPa. Additionally, for each material type, the tests were highly repeatable as seen in the small standard deviations, resulting in high confidence that the printed samples will behave as expected and in the same way every time.

### Puncture and Compliance Testing

The sample size from each section of the heart with the average thickness across the samples at the point of puncture is listed in Table 6 and 7. The inconsistent sample size is due the variability in size of the hearts and the inability to get the proper sample dimensions from every chamber of every heart.

**Table 6:**
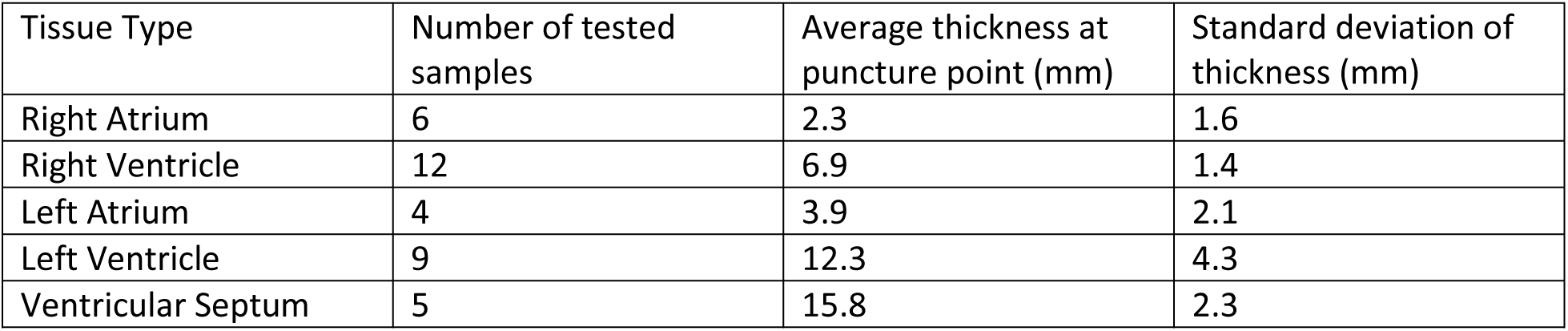
Number of porcine myocardium samples tested with the small pin per tissue type. Thickness at the point of puncture was also measured and averages are recorded here.

**Table 7:** Number of porcine myocardium samples tested with the large pin per tissue type. Thickness at the point of puncture was also measured and averages are recorded here.

| Tissue Type | Number of tested samples | Average thickness at puncture point (mm) | Standard deviation of thickness (mm) |
| --- | --- | --- | --- |
| Right Atrium | 5 | 1.8 | 0.8 |
| Right Ventricle | 11 | 7.3 | 2.5 |
| Left Atrium | 5 | 3.0 | 2.5 |
| Left Ventricle | 7 | 13.1 | 2.3 |
| Ventricular Septum | 5 | 15.1 | 3.4 |

Both Digital Anatomy myocardium and porcine myocardium had similar failure mechanisms. Both saw an initial peak force as the first tough layer was punctured, endocardium for the tissue and the first Agilus layer for Digital Anatomy. The pin then pushed through the myocardium and printed infill. Before the final failure, the epicardium and second Agilus layer delaminated and tented away from the rest of the sample before the pin popped completely through the sample. The delamination is shown in Figure 8. The puncture process for Digital Anatomy is shown in Figure 9. The stiffer infill values of the Digital Anatomy myocardium produced less delamination. The larger pin produced more delamination across samples.

**Figure 8:**
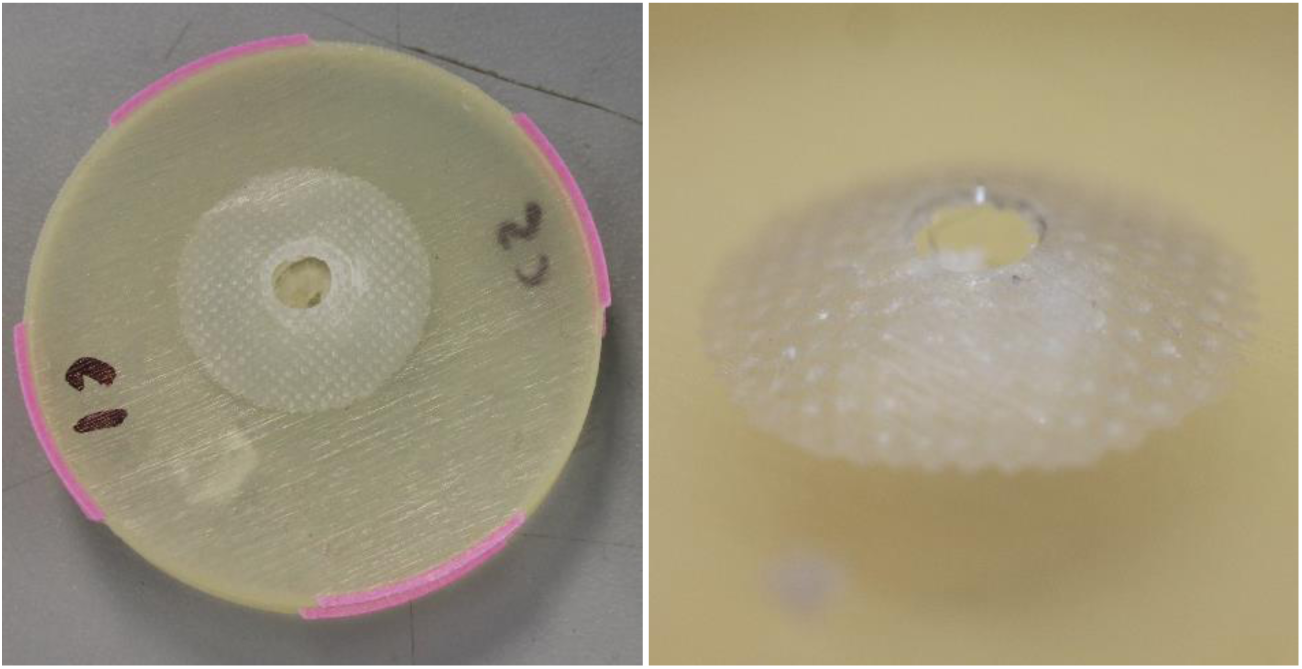
The delamination seen in the puncture test for Digital Anatomy myocardium.

**Figure 9:**
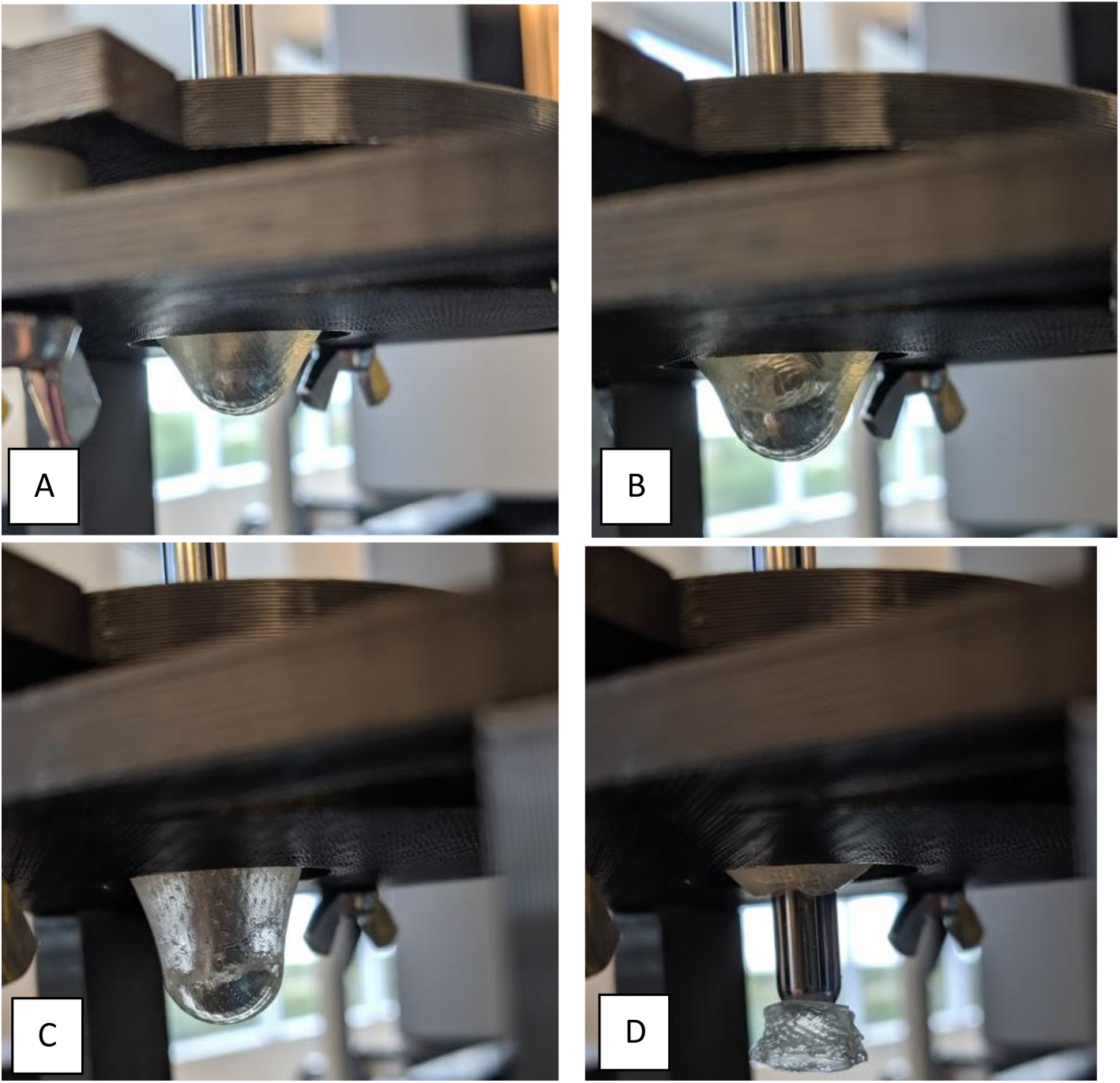
The puncture test for Digital Anatomy Myocardium. (A) Tenting of the entire sample. (B) After puncturing the first layer of Agilus the infill begins to tear. (C) The delamination of the bottom layer of Agilus. (D) Complete puncture.

Analysis was done on printed materials and porcine myocardium puncture data to assess differences in compliance. Stiffness values (N/mm) are reported for this test. The lower the stiffness, the more compliant the samples are.

### Porcine Myocardium

In testing hearts from both locations, the decision was made to combine all the data because there appeared to be no significant difference between butcher and preclinical research hearts despite the differences in harvesting and storing methods. Figure 10 shows load versus displacement data that was collected during puncture testing, where a rounded pin was forced through a suspended tissue sample until complete failure. As seen in Figure 10, the data obtained was highly variable with lots of noise from the internal structures of the myocardium. The data from the two sources falls within the same uncertainty ranges, allowing all the data to be pooled together.

**Figure 10:**
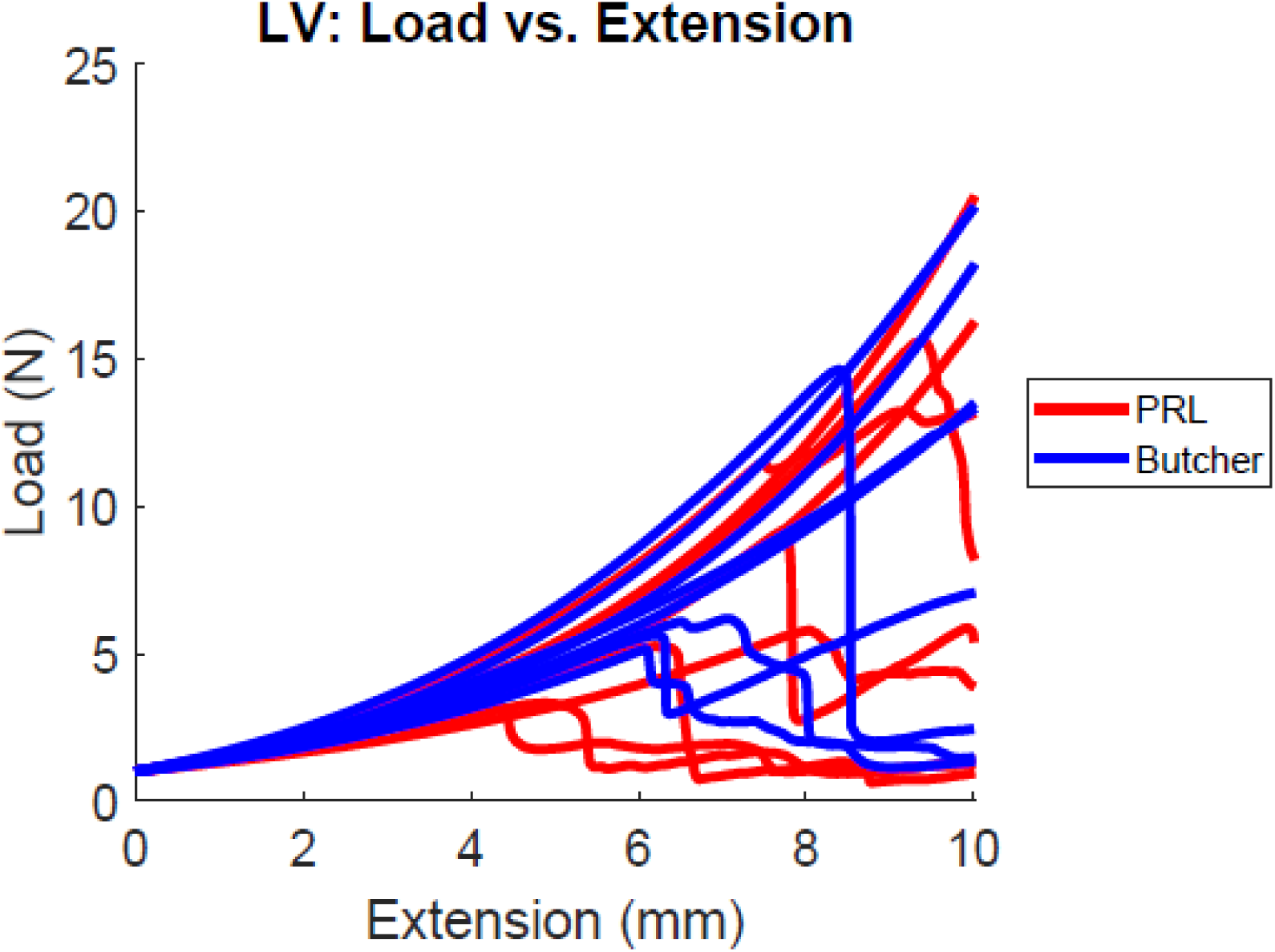
Puncture comparison between butcher hearts and hearts obtained from a pre-clinical research facility.

Stiffness values for two different levels of displacement and pin sizes are shown in Figure 11 and Figure 12. The tissue had significant variability between samples, and more variability using the large pin. The small pin most likely could move around the trabeculae, whereas the large pin had to puncture through all the extra structures. This could be why the large pin has more variability. The small pin also punctured more tissue samples before reaching 10 mm displacement. Once the endocardium is punctured, the use case is no longer typical, so these samples were excluded and resulted in smaller sample sizes for many of the regions. In general, the stiffness increases with displacement and increase in pin size.

**Figure 11:**
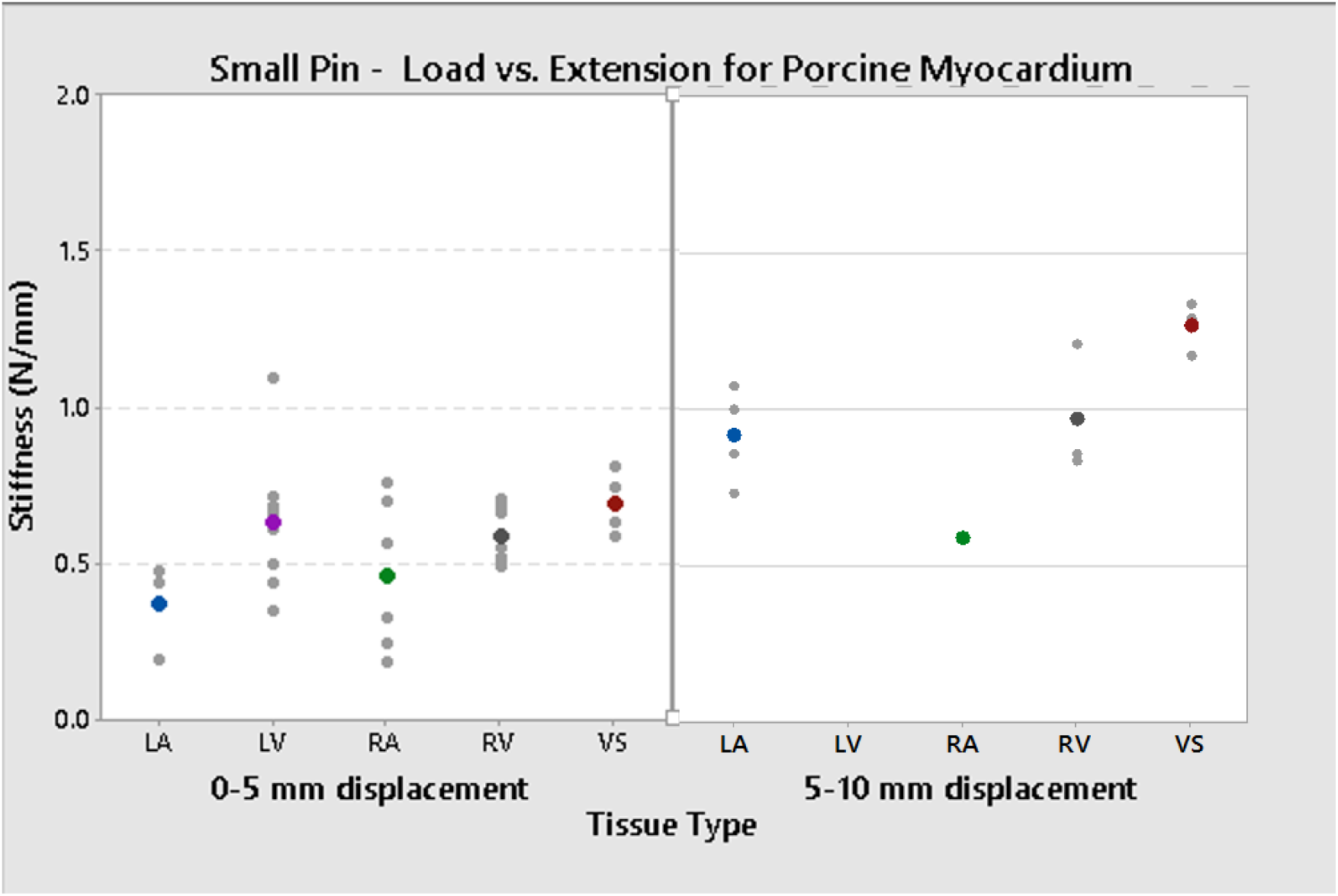
Average slope values and their confidence intervals for two levels of displacement for porcine myocardium from the right atrium, left atrium, right ventricle, left ventricle, and ventricular septum using a small pin.

**Figure 12:**
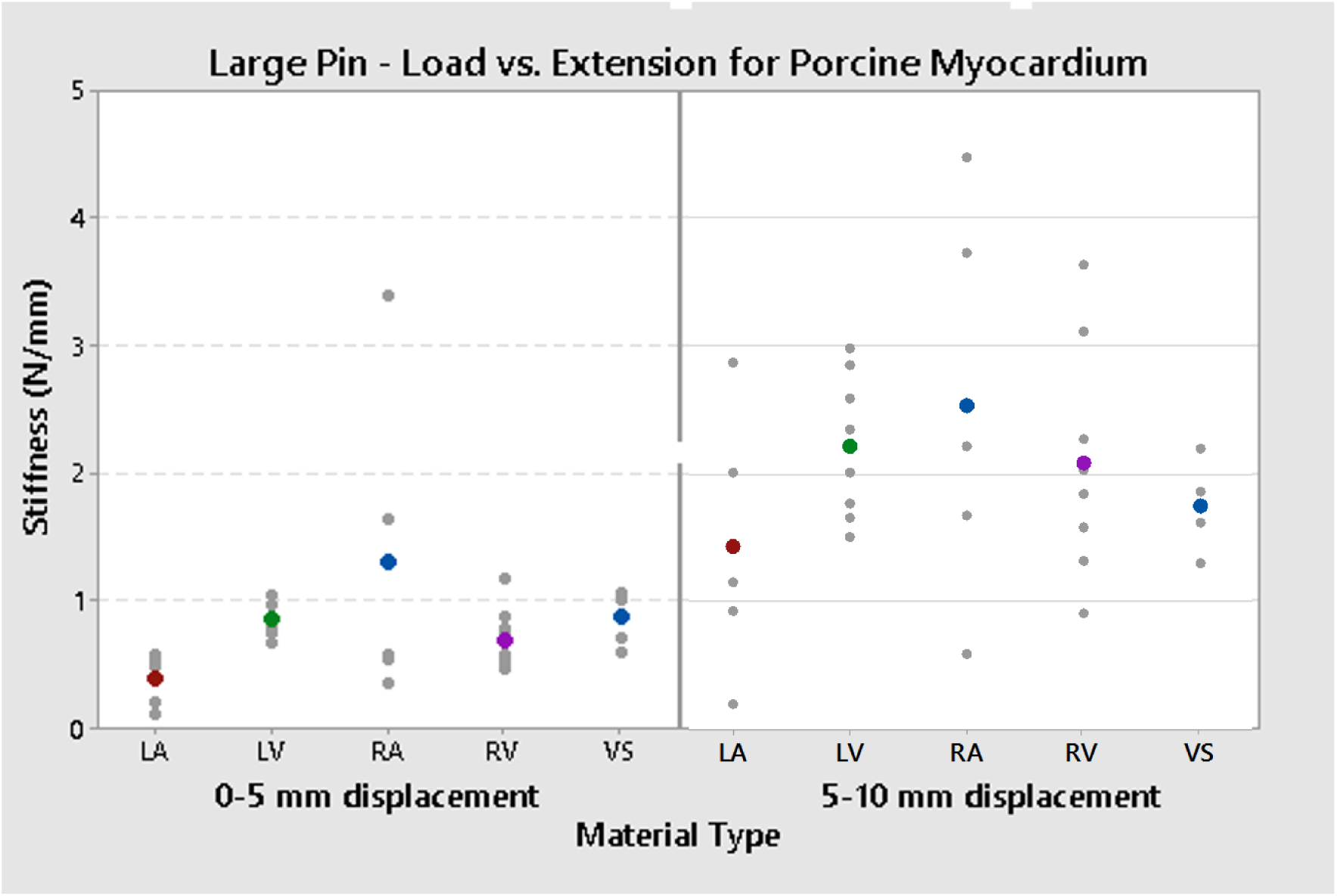
Average slope values and their confidence intervals for two levels of displacement for porcine myocardium from the right atrium, left atrium, right ventricle, left ventricle, and ventricular septum using a large pin.

### Digital Anatomy Printed Material Testing

Compliance data for samples tested at relevant thicknesses is shown in Figure 13 and Figure 14. The printed materials showed very little variability. Increasing the sample thickness from 2.5 mm to 6.6 mm did not seem to affect the compliance for 0-5 mm displacement using either pin size. Using the large pin, changing the thickness from 2.5 mm to 6.6 mm also did not seem to affect the compliance for 5-10 mm displacement. In all cases, the 12.1 mm thick sample was the stiffest. Depending on the use case, adjusting thickness could help achieve targeted stiffness values.

**Figure 13:**
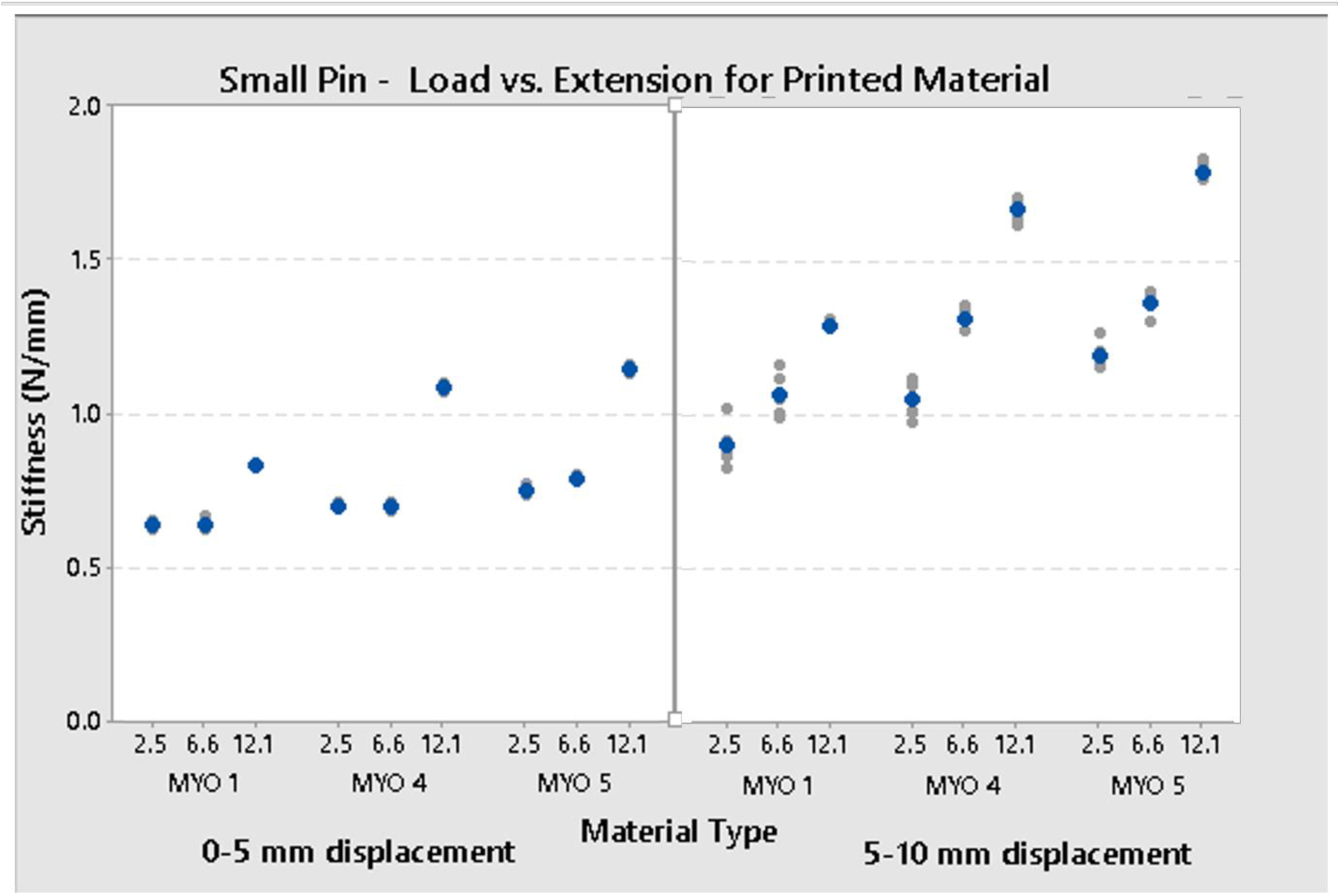
Stiffness values at two displacement levels using a small pin. Three printed digital anatomy materials were tested at three clinically relevant thicknesses.

**Figure 14:**
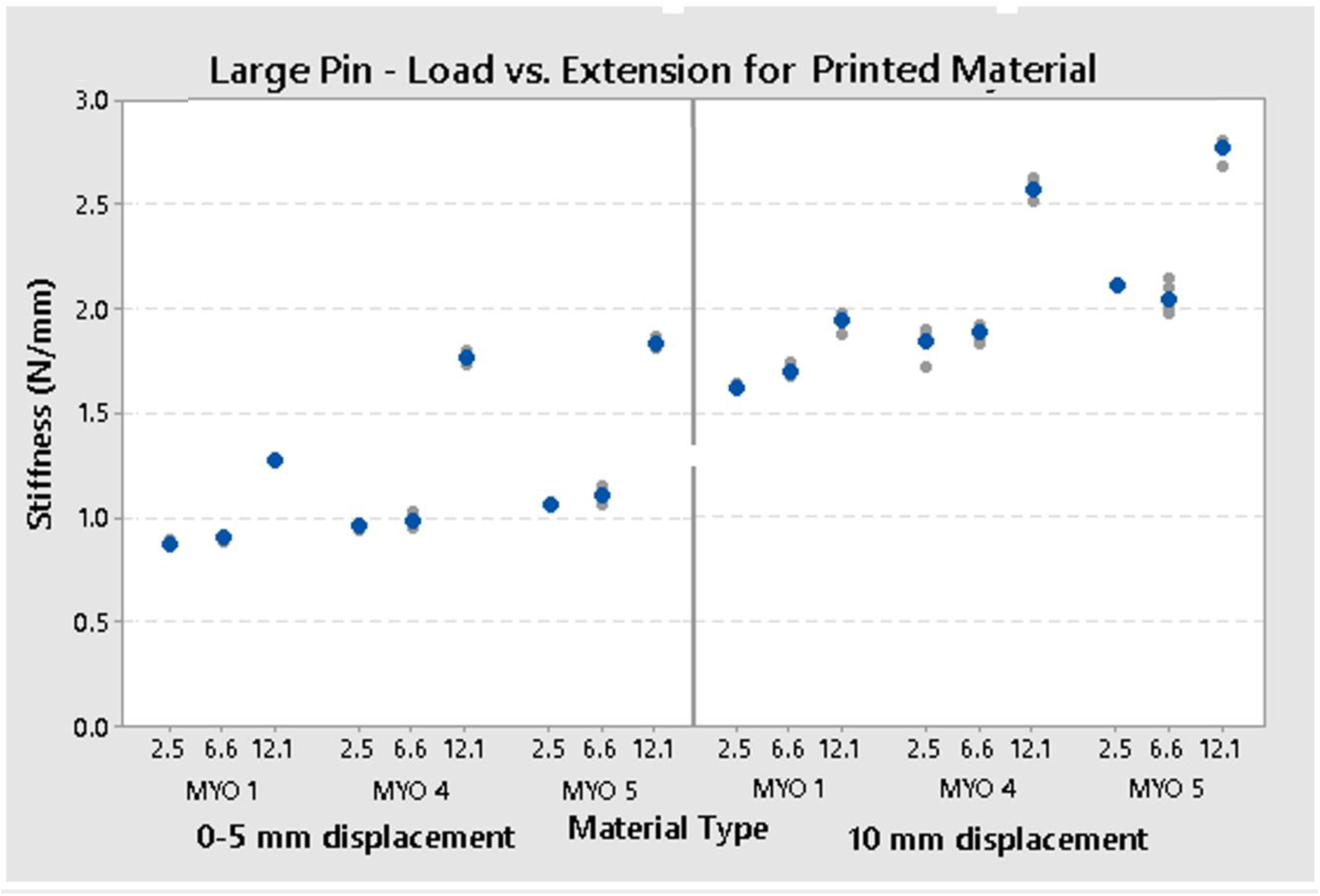
Stiffness values at two displacement levels using a large pin. Three printed digital anatomy materials were tested at three clinically relevant thicknesses.

Changing material type also changed the stiffness values. Myocardium1 is the most compliant and Myocardium 5 is the stiffest. However, for the thinner samples and small levels of displacement the difference in stiffness is small, especially with the more concentrated stress of the smaller pin. Material differences are more apparent with the thicker samples. Given the results of the screening test, it can be assumed that Myocardium 2 and 3 would be between the stiffness values of Myocardium 1 and 4.

### Comparisons Between Printed Materials and Porcine Tissue

Using a small pin at five mm displacement, printed myocardium is much stiffer than porcine myocardium as seen in Figure 15. For ten mm displacement as seen in Figure 16, the printed myocardium falls within the variability of most tissue types. The septum averaged 15.8 mm thickness, and matches well with 12.3 mm thick Myocardium 1. Right ventricle averaged 6.9 mm thickness and the stiffness values for 6.6 mm thickness for all printed myocardium fall within the variability of the right ventricle. However, the average value corresponds best to Myocardium 1. In the tested configurations, the left atrium was too soft to correspond to the printed materials. The right atrium averaged 3.9 mm thickness and the variability encapsulates both the 2.5 mm and 6.6 mm thick Myocardium 1 samples. The 2.5 mm thick Myocardium 4 also corresponds to right atrial stiffness, which implied relevance for Myocardium 2 and 3 at the proper thickness. These results indicate that with the anatomically relevant thicknesses, stiffness values of printed samples can correspond to most of the chambers of the porcine heart.

**Figure 15:**
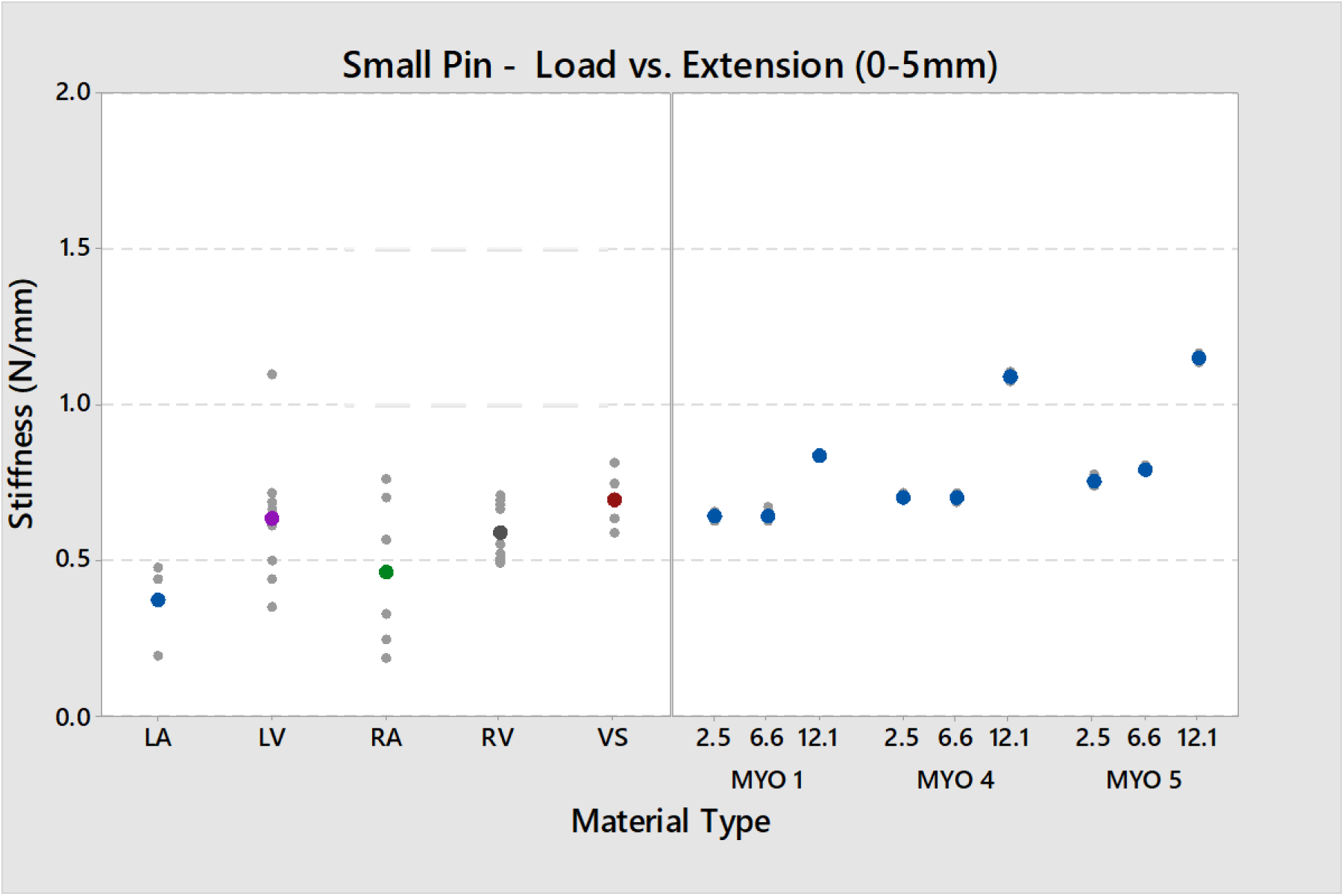
Comparison between porcine myocardium and printed myocardium using a small pin and five millimeters of displacement. The sample size used for porcine myocardium is included on the key. All the printed materials used a sample size of six.

**Figure 16:**
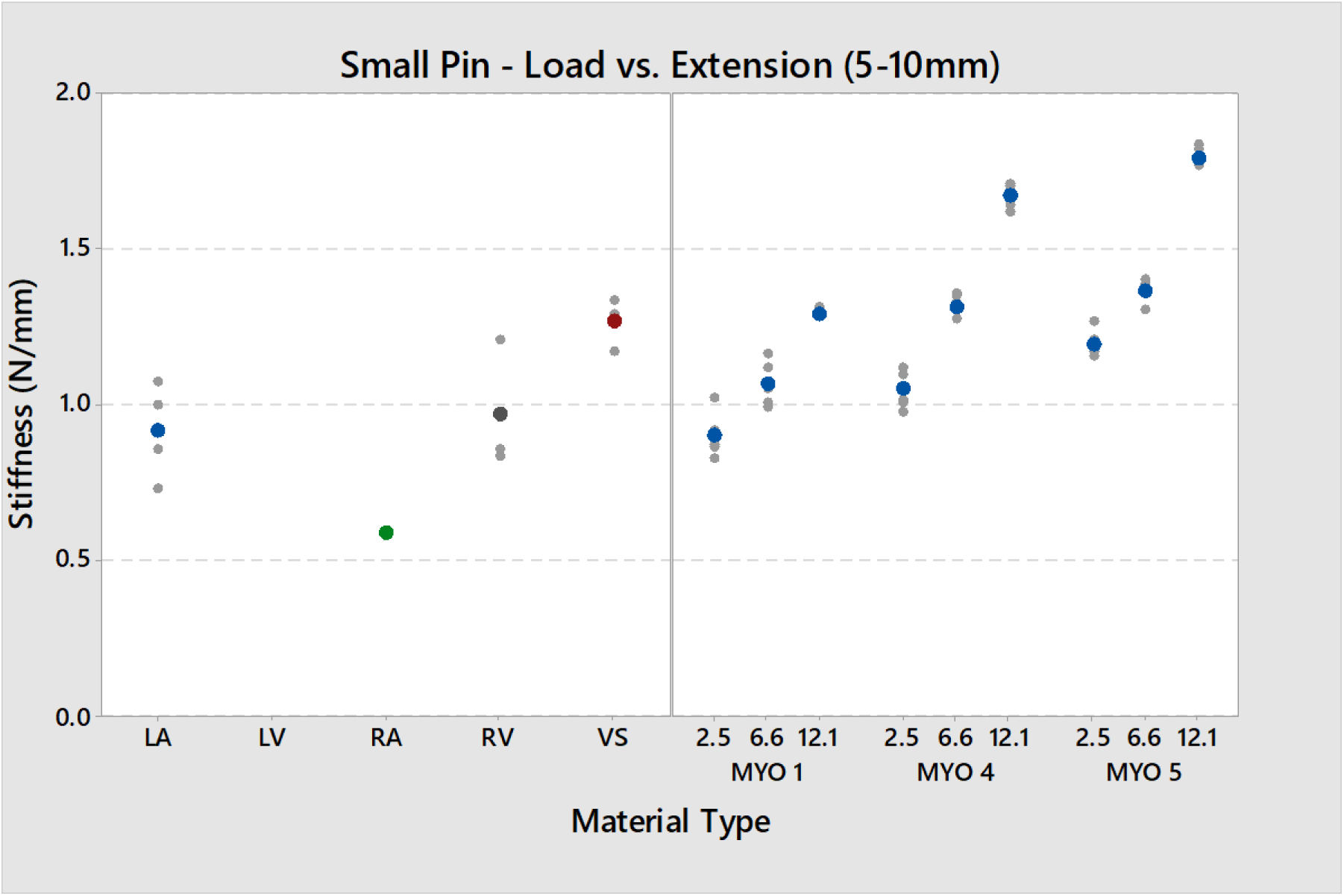
Comparison between porcine myocardium and printed myocardium using a small pin and ten millimeters of displacement. The sample size used for porcine myocardium is included on the key. All the printed materials used a sample size of six.

Using a larger pin, porcine stiffness values still correspond to printed myocardium values, but the relationships do not correspond as well with the matching thicknesses. For example, the left ventricle averaged 13.1 mm thickness, but with five mm displacement the stiffness corresponds best to that of 2.5 mm thick Myocardium 1. For ten millimeters of displacement, left ventricular stiffness could correspond to 12.1 mm thick Myocardium 1, all tested thicknesses of Myocardium 4, or 2.5 or 6.6 mm thick Myocardium 5. If trying to replicate a specific cardiac stiffness, it is important to understand the use case. Boundary conditions, sample geometry, and instruments used on the samples all impact stiffness and need to be chosen accordingly. In general, the larger pin gives more options for stiffness for the different tissue types. By having the stress less concentrated, the stiffnesses from the porcine and printed myocardium overlap much more at both displacement levels, even if thicknesses between the two do not always correspond. The only tissue type that is too soft to match the printed stiffness is left atrium.

**Figure 17:**
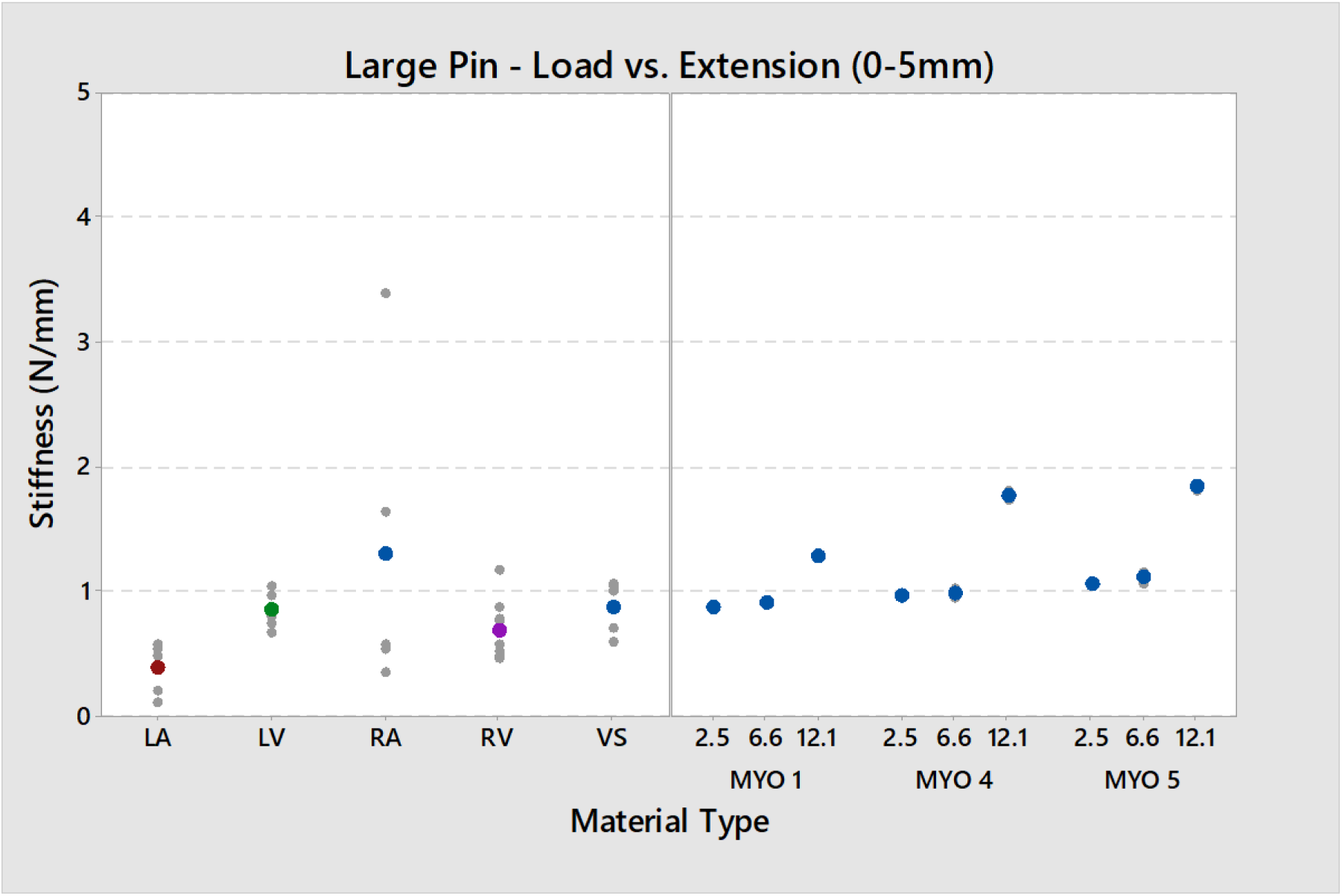
Comparison between porcine myocardium and printed myocardium using a large pin and five millimeters of displacement. The sample size used for porcine myocardium is included on the key. All the printed materials used a sample size of six.

**Figure 18:**
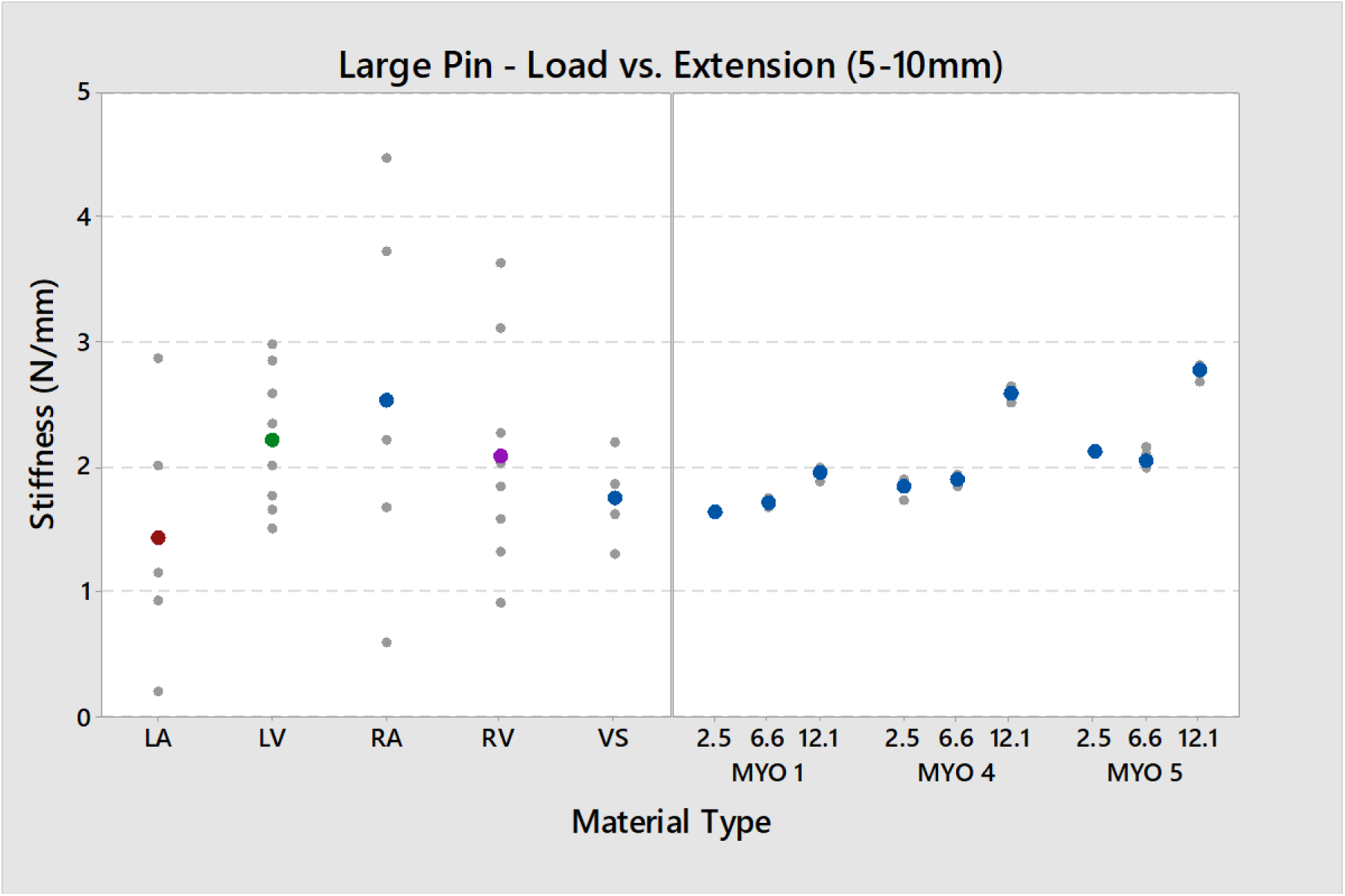
Comparison between porcine myocardium and printed myocardium using a large pin and ten millimeters of displacement. The sample size used for porcine myocardium is included on the key. All the printed materials used a sample size of six.

In addition to similar stiffness values between porcine and printed myocardium, the curves look similar. Figure 19 shows the force vs. displacement curves up to ten millimeters for 6.6 mm thick printed myocardium and right ventricle. For a small pin, the right ventricle thickness averaged 6.9 mm and for large pin it averaged 7.3 mm thick.

**Figure 19:**
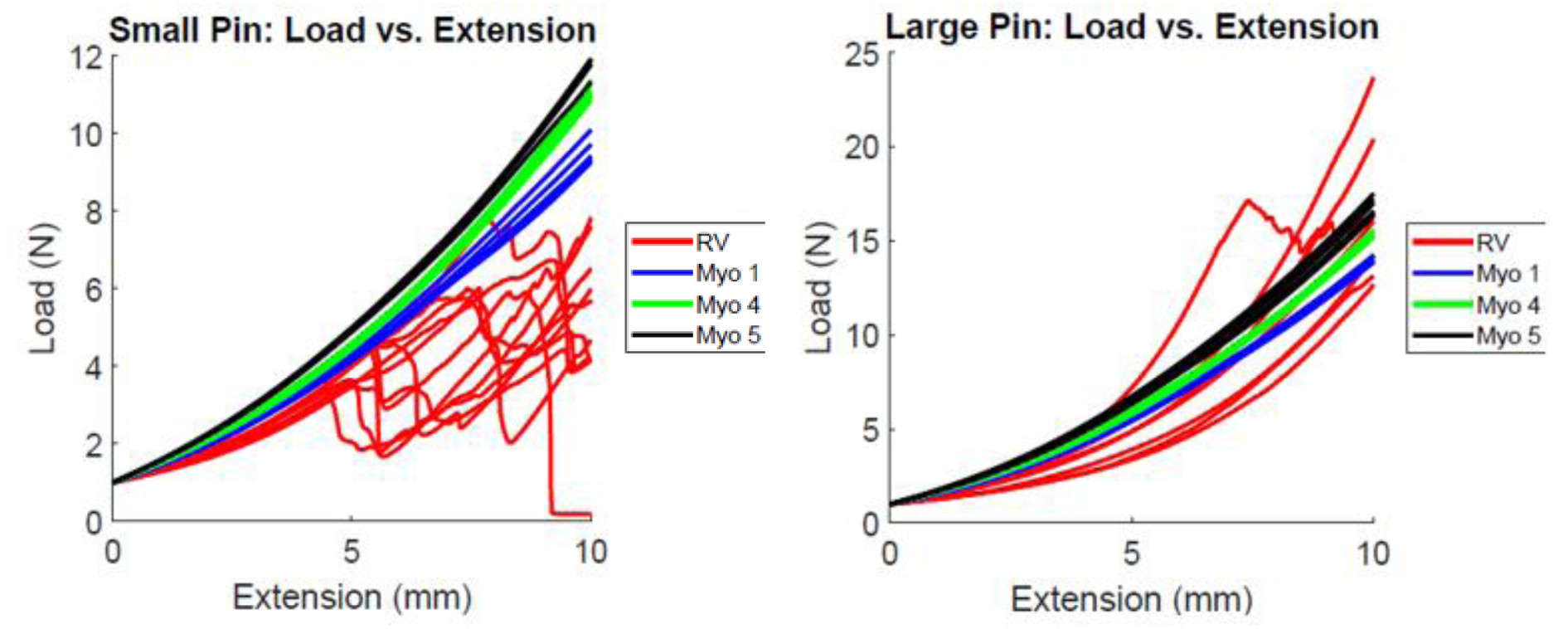
Comparison of the actual force vs. displacement curves for porcine right ventricle and 6.6 mm thick printed myocardium.

Overall, the printed samples show less variability than animal tissue. In general, the first cycle produced slightly higher loads for the displacement compared to the following cycles, but the following cycles overlapped each other as shown in Figure 20. For quantitative curve comparison, the five millimeter displacement slopes were calculated to find the percent differences between cycle curves as shown in Figure 21. The thicker samples showed less percent difference between cycles. All samples showed a noticeable difference between the first cycle and subsequent cycle, suggesting that some pre-conditioning or standard recovery time may be appropriate in order for the parts to fully return to their shape between uses.

**Figure 20:**
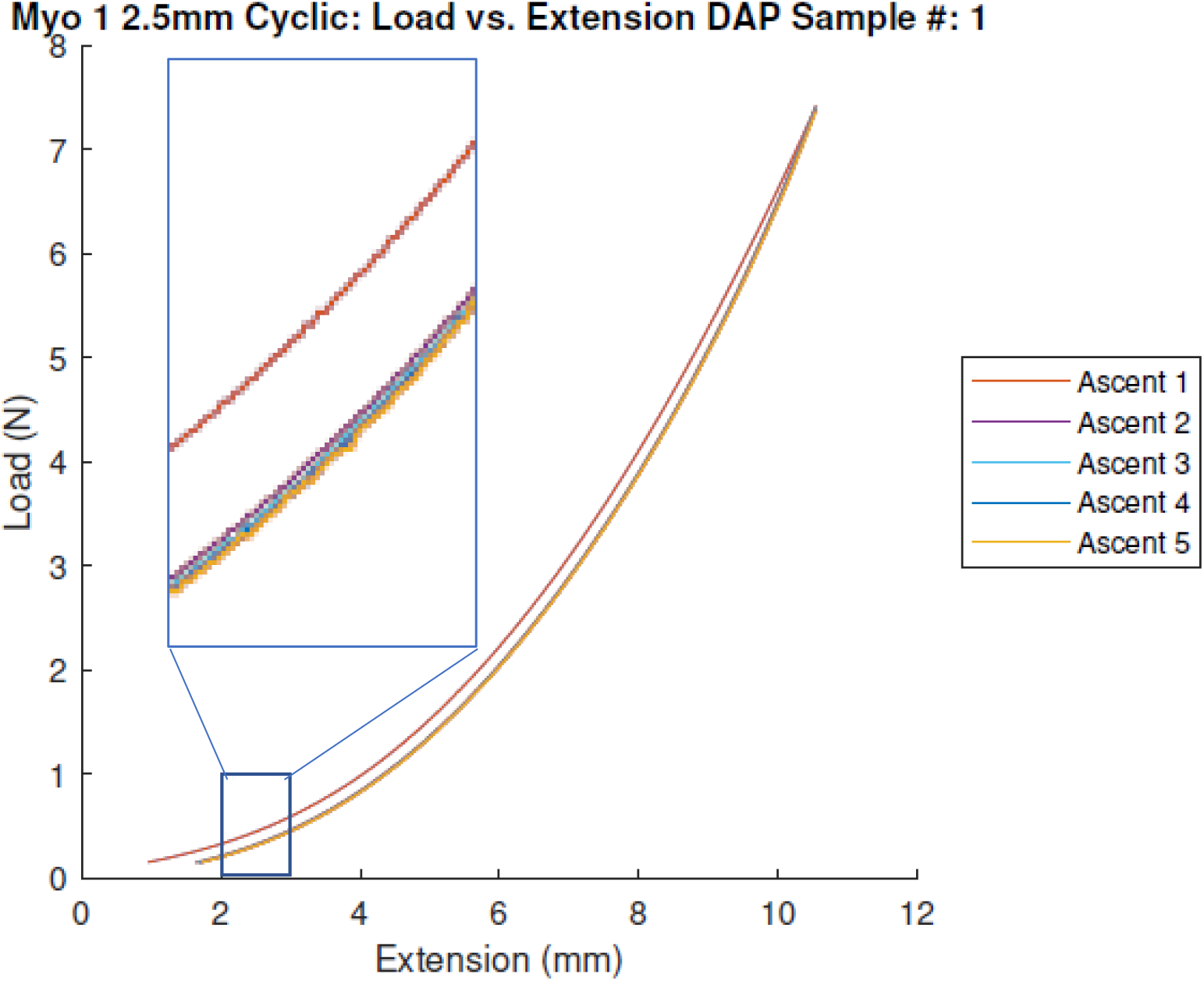
An example of the cyclic compliance testing done on a 2.5 mm thick sample of Myocardium 1. Ascent 1 is the top curve, and Ascent 2-5 overlap each other on the bottom curve.

**Figure 21:**
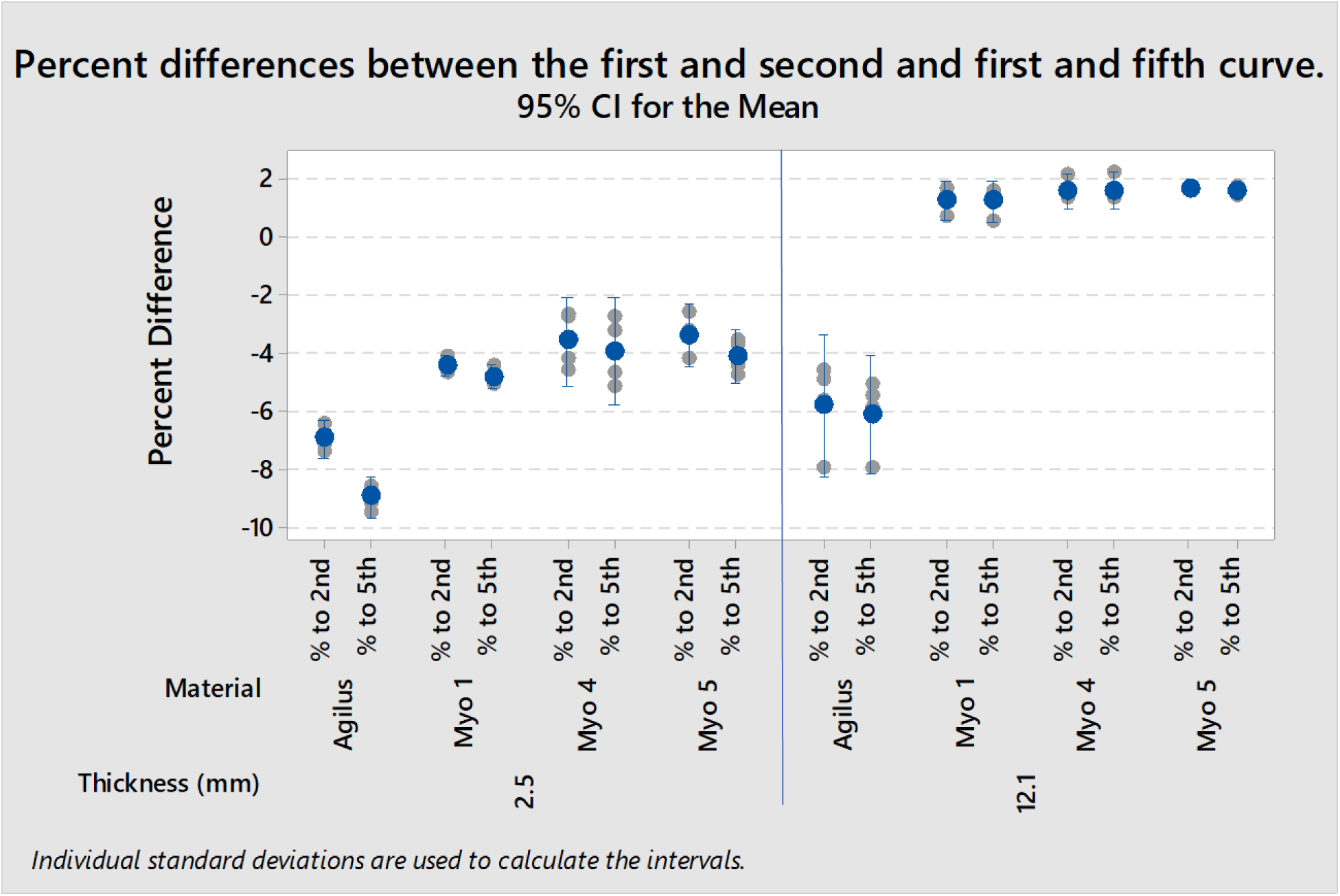
An example of the percent difference analysis using the lead quadratic coefficient for 2.5 mm Myocardium 1 for all four samples.

### Suture Testing

The sutures cut through the printed material more easily than porcine tissue. Average peak suture pull forces are listed in Table 8 and shown in Figure 22. Overall, tissue is 2.4-9 times stronger than printed materials. However, this is also given the specific suture setup used to prevent preliminary tearing of printed samples. Standard suture and tie techniques would result the printed material failing prior to testing. Regardless, the tissue and printed myocardium both showed delamination prior to failure as shown in Figure 23.

**Table 8:** Average peak suture pull forces for porcine myocardium and printed materials.

| Material | Average peak suture pull force (N) |
| --- | --- |
| 12 mm Agilus | 4.754 |
| 4 mm Agilus | 4.239 |
| 12 mm Myo 1 | 1.663 |
| 4 mm Myo 1 | 1.765 |
| 12 mm Myo 5 | 3.450 |
| 4 mm Myo 5 | 2.881 |
| RV | 16.469 |
| LV | 11.387 |

**Figure 22:**
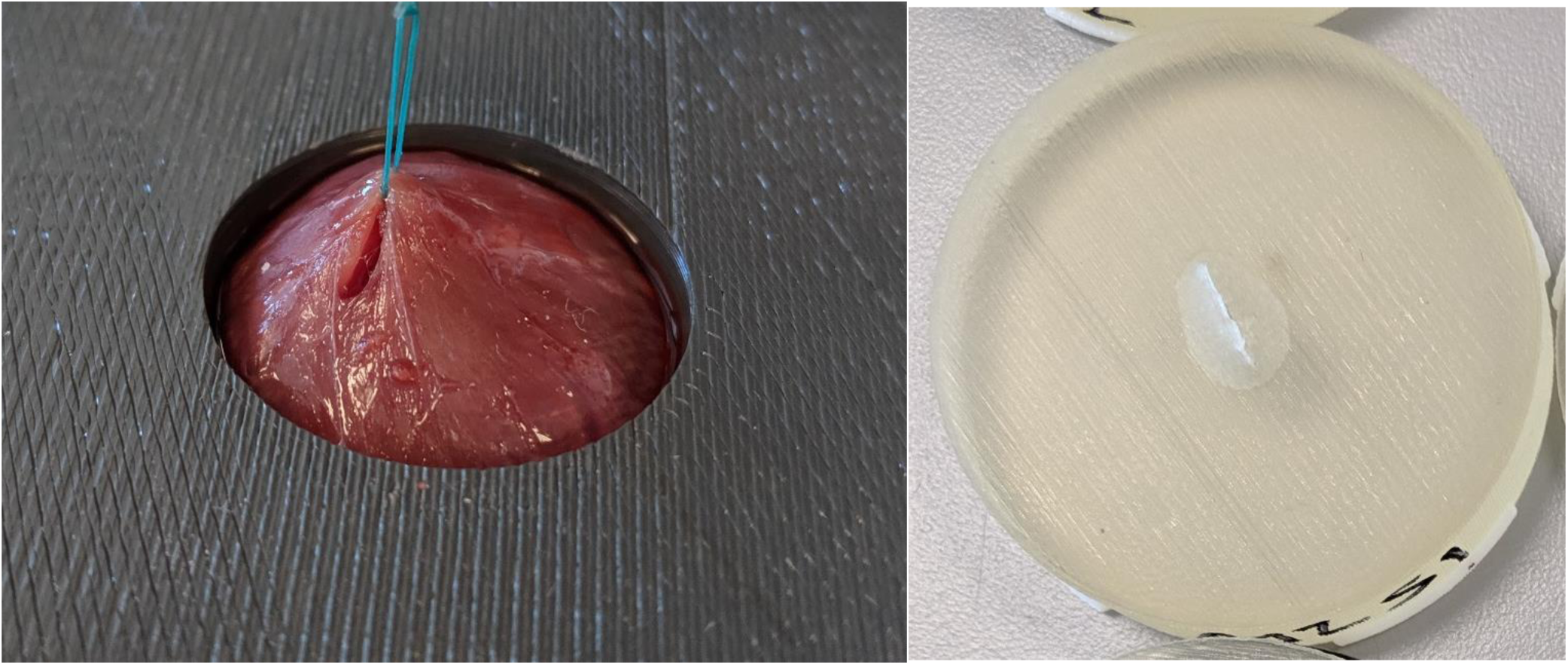
(Left) Porcine myocardium failing during the suture test. Note the hole forming as the suture pulls on the epicardium. (Right) Myocardium 1 after the suture pull test. The bubble around the cut shows the delamination of Agilus before failure.

**Figure 23:**
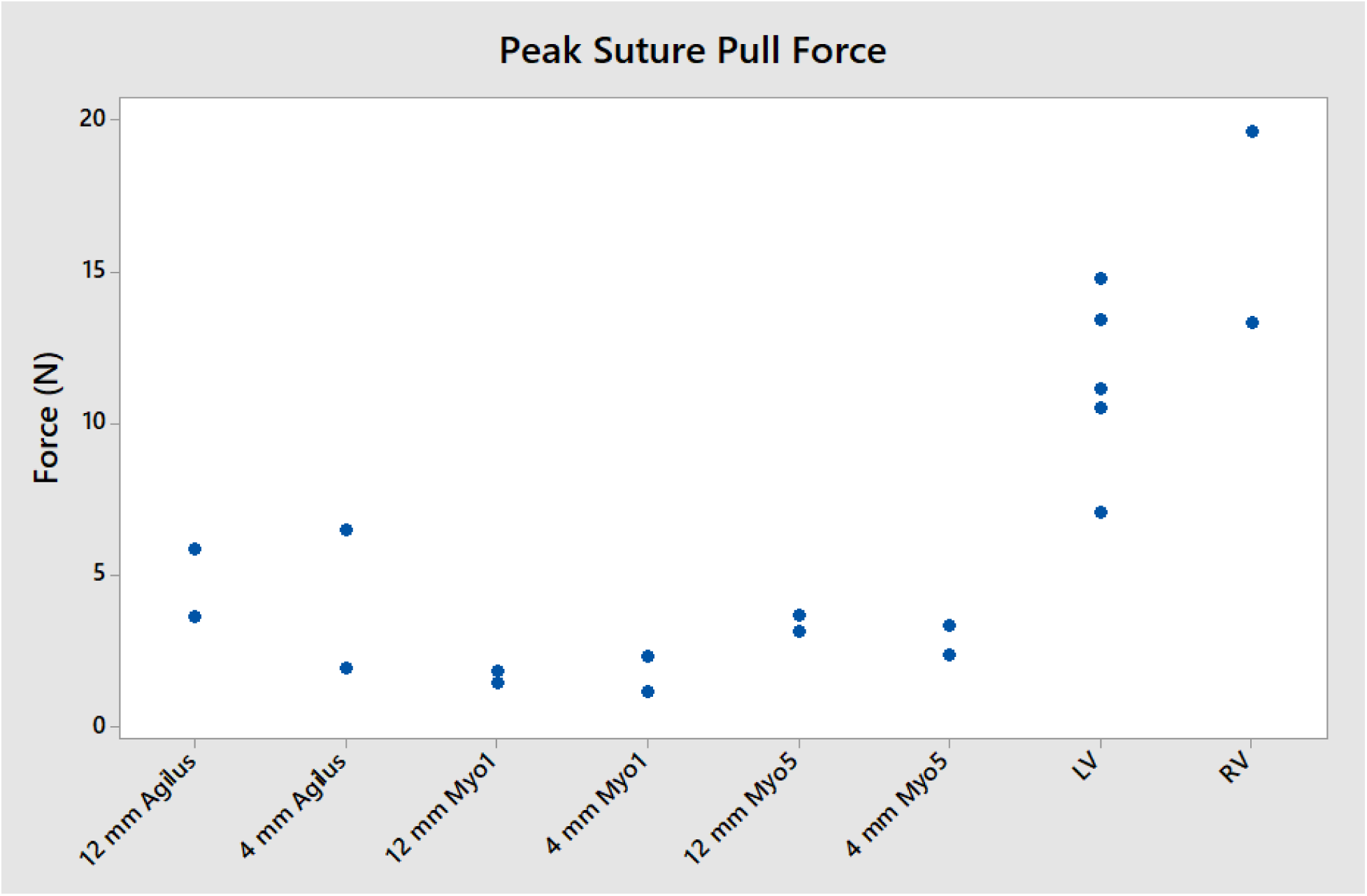
Peak Suture Pull force for printed samples and porcine myocardium.

### Qualitative Assessment for cutting and suturing

Overall, the reviewers thought the printed myocardium was much closer to real tissue than Agilus, but still inferior, as seen in the scores in Table. For suturing, the printed myocardium crumbled and tore too easily. The Agilus layer tented before puncturing of the suture needle, requiring a greater initial force. The drag felt on the needle when moving through the material was judged to be too high. When tightening the knot, the suture tore through the printed myocardium easily, preventing use in any appropriate suturing applications. For cutting purposes, the printed myocardium was too sticky and required more force than cutting actual myocardium.

**Table 9:** Scores comparing printed materials to porcine myocardium for cutting and suturing. The assessment was performed by four individuals with years of animal and cadaver pre-clinical research experience. The scores range from 1 (not at all similar) to 7 (very similar to tissue).

| Reviewer | Suture Agilus | Suture Myo1 | Suture Myo5 | Cut Agilus | Cut Myo1 | Cut Myo5 |
| --- | --- | --- | --- | --- | --- | --- |
| A | 1 | 2 | 6 | 1 | 6 | 2 |
| B | 1 | 3 | 5 | 1 | 3 | 3 |
| C | 1 | 2 | 2 | 1 | 1 | 1 |
| D | 1 | 3 | 6 | 1 | 3 | 2 |

The reviewers thought the printed myocardium needed to be more compliant, more lubricious, and tougher in order to mimic real myocardium. They said the feel of the Digital Anatomy was getting close to real myocardium but was lacking some compliance. They also thought better lubricity would reduce the additional drag experienced when suturing and cutting, and they mentioned that materials need to be tougher in order to accommodate more realistic suture techniques and holding forces. However, for applications that do not require cutting or suturing they thought the compliance of Digital Anatomy could be a useful resource for preliminary bench testing.

## Conclusions

The 3D printed myocardium shows promise with its ability to simulate porcine tissue compliance. While for some use cases the printed myocardium appears to be too stiff, specifically for smaller displacements with higher concentrated stress, adjustments could be made to match the stiffness of the targeted tissue type in terms of material composition, geometry, boundary conditions, and use case. For example, for some use cases in order to match the stiffness of the left ventricle the thickness of a sample printed with Myocardium 1 can be reduced by about half.

The biggest difference between the porcine and printed myocardium was the variability. As previously discussed, the printed materials are highly consistent while the tissue is highly variable. The tissue has trabeculae and muscle fibers that are unique to each heart and are a significant contributing factor to the tissue variability. Even though the printed myocardium may not match the tissue stiffness in every use case, the repeatability of the materials can be a significant advantage when used in bench models for testing.

Furthermore, the porcine and printed myocardium failed in similar ways when they were puncture tested. The porcine myocardium has thin, tough layers of tissue on the inside and outside of the heart, namely the endocardium and epicardium. Similarly, the printed myocardium is wrapped in a thin layer of Agilus. During the puncture test, both samples saw the same pattern before reaching failure. The first tough layer would tent into the sample before breaking and creating the first large peak on the force vs. displacement graph. Then the pin would move through the center of the sample, tent and puncture the second tough layer of the sample. This same puncture profile was also seen in the suture pull force testing, where the tough layer would pull up and delaminate from the center before the suture cut through it. While it is encouraging that the printed myocardium is failing in the same way as porcine myocardium, in general the porcine myocardium is still both stronger and more compliant.

Additionally, printed specimens representing myocardium are a beneficial development tool. In development work, especially when comparing the function of different tools, repeatability between samples and times of testing is very important to minimize confounding variables. Digital Anatomy printed myocardium shows high repeatability in stiffness value within the same sample tested multiple times, as well as between samples. This is expected to allow a bench model printed with these materials to perform repeatably over the course of a test, as well as perform repeatably between similar bench models. Additionally, they show less part to part variation than porcine myocardium and better cyclic repeatability than Agilus.

For the cases where bench models need to have a more realistic feel, printed myocardium is better than Agilus. Agilus is much less compliant than the printed myocardium and unrealistically stiff compared to porcine myocardium. Even though suturing and cutting through the printed samples have major limitations compared to porcine myocardium, Agilus performed worse than the printed myocardium materials.

Despite the limitations of the Digital Anatomy materials to simulate cutting and suturing, the materials show good promise when it comes to repeatability and compliance related to porcine tissue. This allows for the materials to be used to create bench models for device testing, as well as anatomical models to simulate procedures for development and training.

